# Epithelial-Mesenchymal Wnt Crosstalk Directs Planar Cell Polarity in the Developing Cochlea

**DOI:** 10.64898/2026.02.24.705389

**Authors:** Ippei Kishimoto, Abel P. David, Kevin P. Rose, Balasubramanian Narasimhan, Bradley Efron, Sara E. Billings, Erin L. Su, Wuxing Dong, Taha A. Jan, Ronna Hertzano, Alan G. Cheng

**Author notes:** Corresponding author Alan G. Cheng, M.D. 801 Welch Road Department of Otolaryngology-HNS Stanford, CA 94305.

## Abstract

Cochlear hair cells are crucial for sound transduction and are precisely oriented by planar cell polarity (PCP). Wnt proteins direct PCP in vertebrates, but their roles in regulating cochlear PCP remain unclear. Here, we inhibited Wnt secretion by ablating Wntless in the cochlear epithelium and found shortened cochlea, mild hair cell misorientation and mislocalization of PCP proteins, Fzd6 and Dvl2, representing defects less severe than classic PCP mutants. Computational cell-communication analysis predicted that candidate Wnts from the cochlear epithelium (Wnt5a, Wnt7a, Wnt7b) and surrounding periotic mesenchyme (Wnt5a) act on developing hair cells. Deletion of Wnt5a, Wnt7a, and Wnt7b additively shortened the cochlea without causing hair cell misorientation or mislocalization of PCP proteins. Moreover, deletion of Wnt5a alone in the periotic mesenchyme failed to cause PCP defects. However, ablating both epithelial and mesenchymal Wnts caused a severely shortened cochlea with apically malrotated hair cells devoid of polarized core PCP proteins. Thus, Wnts serve as global instructive cues directing cochlear outgrowth and hair cell polarization, with remarkable redundancy of distinct Wnts across epithelial and mesenchymal compartments to ensure a fail-safe developmental program.

## Introduction

The cochlea is a spiral-shaped sensory organ critical for sound transduction. As mechanotransducers of sound vibration, hair cells are precisely arranged both tonotopically along the length of the cochlea and radially to form three rows of outer hair cells (OHCs) and one row of inner hair cells (IHCs).

During embryonic development in mice, the cochlear duct initially forms as an outpouching from the ventral otocyst at embryonic day (E) 11 and subsequently elongates to become a two-and-a-half-turn organ^1, 2^. In the floor of the cochlear duct, terminal mitosis of prosensory cells begins in the apex and extends towards the base, with hair cell specification occurring in the opposite direction between E12.5 and E14.5^2, 3, 4^. Four rows of hair cells and interdigitating supporting cells are arranged in a checkerboard pattern and a precise radial orientation, ensuring that hair bundles, which house the mechanotransduction machinery, uniformly face laterally. This organized cochlear patterning is regulated by the planar cell polarity (PCP) pathway.

The PCP pathway is evolutionarily conserved from Drosophila to mammals, governing both convergent extension and hair cell polarity in the developing mouse cochlea^5, 6^. In humans, mutations underlying Bardet–Biedl syndrome^7^ and Alstrom syndrome^8, 9^ disrupt PCP signaling and cochlear hair cell development, resulting in hearing loss. In mammals, there are six groups of core PCP proteins-Frizzled (Fzd), Van Gogh (Vangl), Celsr1, Dishevelled (Dvl), Prickle (Pk), and Diego (Dgo)-all of which are asymmetrically expressed at opposite poles of cochlear hair cells and supporting cells^10, 11, 12, 13, 14, 15^. Deletion of Celsr1 or Vangl2 causes cochlear shortening and severe misorientation of hair cells^16, 17^, whereas mutants of Vangl1 displayed relatively mild PCP defects^16^. Concurrent deletion of Fzd3/Fzd6 or Dvl1/Dvl2 leads to striking PCP defects^12, 14^, and deletion of other core PCP proteins in the presence of the Vangl2 looptail mutation amplifies the severity^16^, suggesting complementary and redundant roles among core PCP proteins.

Wnt signaling plays multiple essential roles during development, including cell differentiation, proliferation, cell polarity, and tissue patterning^18, 19^. The Wnt signaling pathway can be broadly classified as canonical and non-canonical, where the former is mediated by β-catenin and the latter by PCP signaling^20^. In the cochlea, Wnt/β-catenin signaling is required for hair cell specification^21^, radial patterning^22^, and supporting cell specification^23^. In vertebrates, although Wnts are known to regulate PCP signaling in various organs such as the neural tube, muscles, and limbs^24, 25^, whether this occurs in the cochlea is unclear.

Among the 19 Wnt ligands in mammals, Wnt2b, Wnt4, Wnt7a, Wnt7b, Wnt9a, and Wnt11 are expressed in the developing cochlear epithelium and Wnt5a is expressed in both the cochlear epithelium and periotic mesenchyme (POM)^15, 26, 27^. Wnt7a mutant cochleae show normal hair cell orientation^28^. While Wnt5a knockout cochleae display defects in convergent extension, hair cell orientation, and localization of Vangl2, although the defects were minor and penetrance was limited, with only one third of mutants affected^29^. We and others have previously shown that concurrent ablation of Wnt members in the cochlear epithelium (by deleting Wntless or Wnt5a and Wnt7b) caused a significant shortening of the cochlea, mild misorientation of hair cells, and mislocalization of a subset of core PCP proteins (Fzd6 and Dvl2) in hair cells^15, 30, 31^. Because the severity of hair cell polarity defects observed in these studies is relatively mild in comparison to other PCP mutants (e.g., Vangl2-looptail^16^), the requirement of Wnts for PCP signaling I the cochlea has remained controversial^32, 33^.

Here, we used computational analysis of single-cell transcriptomic data from the developing cochlea and identified Wnt5a, Wnt7a, and Wnt7b in the cochlear epithelium and Wnt5a in the surrounding POM as the top candidate Wnts that act on developing hair cells. Deletion of multiple Wnts (Wnt5a, Wnt7a, and Wnt7b) from the cochlear epithelium shortened the cochlear duct, without causing misorientation of hair cells or mislocalization of core PCP proteins, suggesting a redundancy among Wnts. Inhibiting the secretion of both epithelial and mesenchymal Wnts further shortened the cochlea and led to severely malrotated hair cells and loss of polarization of all PCP proteins examined, with phenotypes remarkably more severe than Wntless ablation in the cochlear epithelium alone. Collectively, our study indicates that multiple secreted Wnts from both the epithelial and mesenchymal compartments are required for cochlear outgrowth, hair cell orientation and PCP signaling.

## Results

### Ablating Wntless from the cochlear epithelium disrupts cochlear outgrowth and hair cell polarization

Wntless (Wls) is a chaperone transmembrane protein required for secretion of Wnt proteins, a process essential for all Wnts^34, 35^. We found that Wntless mRNA is expressed throughout the developing cochlear epithelium (Fig. 1b, Supplementary Fig. 1c-c”, e-e”, g-g”), corroborating previous results^15^. To ablate Wntless in the cochlear epithelium prior to hair cell differentiation, we first established Emx2-Cre; Wls-flox mice. Emx2-Cre activity starts in the otocyst as early as embryonic (E) 10.5 (Fig. 1a)^36^. In E16.5 *Emx2^Cre/+^; Wls^flox/flox^* (Emx2-Wls cKO) cochleae, there was almost no detectable Wls mRNA in the cochlear epithelium, whereas that in the POM remained comparable to those in control cochleae (Fig. 1c, Supplementary Fig. 1d-d”, f-f”, h- h”), indicating efficient and specific deletion of Wls within the cochlear epithelium in Emx2-Wls cKO mice. At E18.5, Emx2-Wls cKO cochleae are significantly shorter (∼39%) than those from wildtype littermates (2,856±381 and 4,653±220 μm, respectively, p < 0.0001) (Fig. 1d, e). Moreover, Emx2-Wls cKO cochleae displayed significantly fewer (∼29%) Myosin7A-positive hair cells (mean 1,917±234 and 2,692±156 per cochlea, respectively, p < 0.01, Fig. 1f). These findings suggest that inhibiting Wnt secretion in the cochlear epithelium leads to defects in cochlear outgrowth and/or convergent extension. To assess hair cell orientation in phalloidin-stained E18.5 cochleae (Fig. 1g, h, Supplementary Fig. 1i, j), we measured the average and variability of orientation of each row of hair cells (circular mean and variance). In control mice at E18.5, cochlear hair cell bundles face laterally with the most lateral row of hair cells (the third row of OHCs) slightly rotated toward the apex (Fig. 1g-g’, Supplementary Fig. 1i-i’), consistent with previous reports^28, 37^. Compared to controls, Emx2-Wls cKO cochlear hair cells exhibited significantly larger variance in the middle and basal turns (circular variance ratio: 2.00, Supplementary Table 1), along with a mild, but significantly different mean orientation of HCs (IHC, OHC1, and OHC3+, Fig. 1h-h’, i, Supplementary Fig. 1j-j’, k, Supplementary Table 1). We also examined the molecular basis of cochlear PCP by assessing the polarized expression of core PCP proteins in hair cells. Hair cells display a complement of core PCP proteins^15^ (Fig. 1j). In the control cochlea, we found Fzd6, Vangl2, and Celsr1, to be localized to the medial junctions and Dvl2 to the lateral junctions of OHCs (Fig. 1k-k’’’, m, o). By contrast, the Emx2-Wls cKO cochlea lacked polarized expression of Fzd6 and Dvl2 while Celsr1 and Vangl2 remained polarized (Fig. 1l-l’’’, n, p). This finding suggests that Wnt secretion from the cochlear epithelium is required for establishing and/or maintaining the polarization expression of a subset of, but not all, core PCP proteins. Taken together, our data suggest that disruption of Wnt secretion from the cochlear epithelium prevents elongation of the organ and causes a mild PCP defect (shortened cochlea, hair cell misorientation, and loss of polarized expression of a subset of PCP proteins), corroborating previous reports^15, 31^.

**Figure 1.**
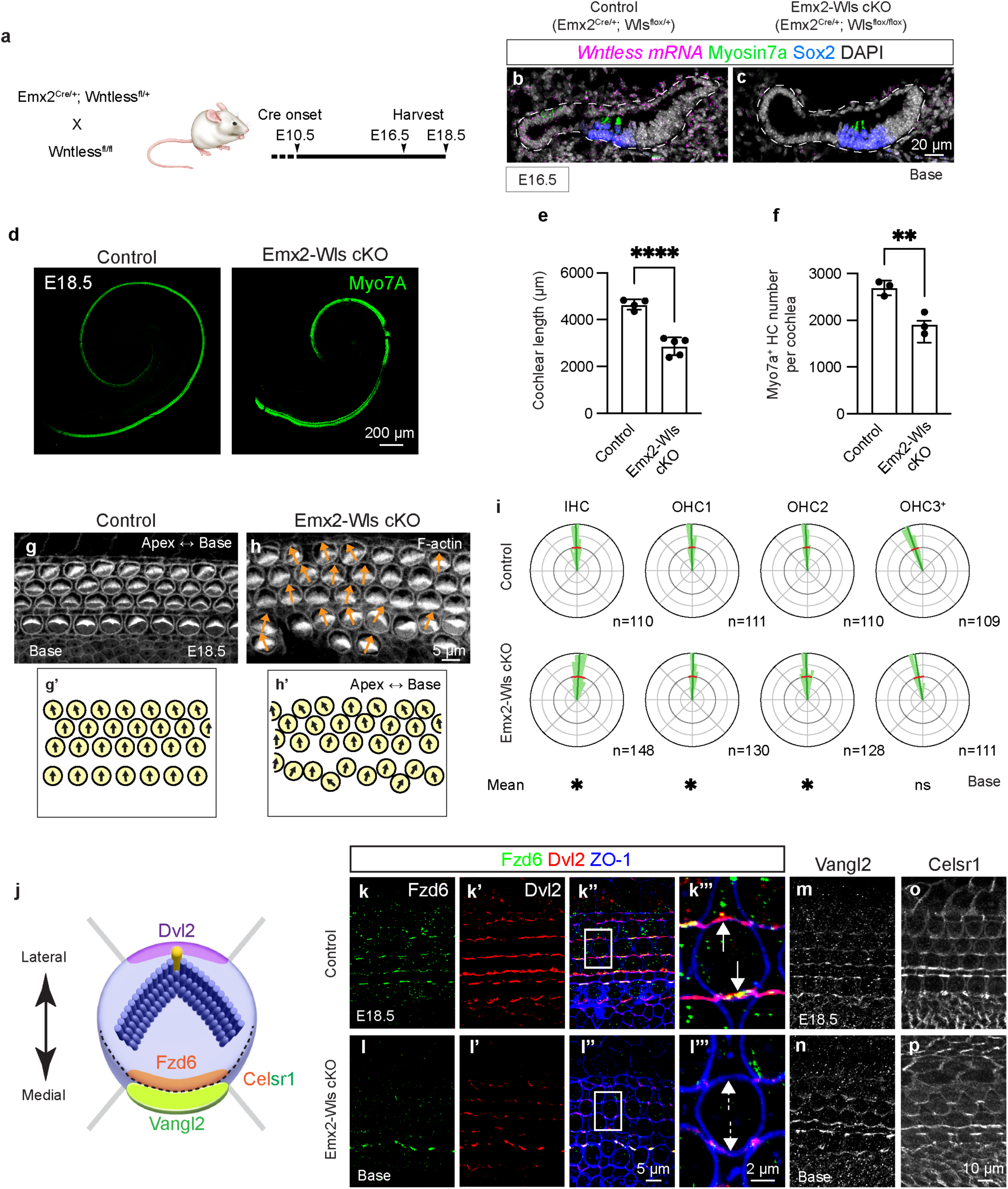
Deletion of Wntless shortens the cochlea and causes mild hair cell malrotation. **a.** Transgenic approach to delete Wntless in the cochlear epithelium. Cre recombinase in the Emx2-Cre cochlear epithelium begins around embryonic day (E) 10.5. **b-c.** Immunostaining and fluorescence *in situ* hybridization showing *Wls* mRNA in the cochlear epithelium in controls (*Emx2^Cre/+^; Wls^flox/+^*), and no Wls mRNA signals in the cochlear epithelium of *Emx2^Cre/+^; Wls^flox/flox^* (Emx2-Wls cKO) mice. **d.** Whole mount preparation of E18.5 cochlea immunostained for Myo7A, showing shortened Emx2-Wls cKO cochlea. **e.** Emx2-Wls cochleae are significantly shorter than controls at E18.5. **f.** There are fewer Myo7A-positive hair cells per cochlea in Emx2-Wls cKO than controls. **g-h.** Hair cells in Emx2-Wls cKO cochlea exhibit more variable orientations compared to controls. **i.** Circular histograms depicting the distribution of hair bundle orientation in the basal turn of control and Emx2-Wls cKO cochleae, with the latter showing significantly larger circular variance and different average orientation in the IHC, OHC1, and OHC3+ rows. The OHC3+ row represents the third and more lateral rows of OHCs. **j.** Cartoon illustrating the asymmetric localization of core PCP proteins in hair cells. Dvl2 is present at the lateral pole of HCs, while Fzd6 is at the medial pole. Vangl1 and Vangl2 are present at the lateral side of SCs, and Celsr1 is enriched at the junction between the medial side of HCs and the lateral side of SCs. **k-l.** Immunostaining for Fzd6 and Dvl2, and ZO1 in the basal turn of cochleae from control (k-k’’’) and Emx2-Wls cKO (l-l’’’) mice. Polarized expression of Fzd6 and Dvl2 is observed at the medial and lateral poles of OHCs, respectively, in the control (k’’’, white arrows) but not in the Emx2-Wls cKO cochlea (l’’’, dashed white arrows). **m-p.** Immunostaining showing expression of Vangl2 and Celsr1 in the medial pole of hair cells in control (m, o) and Emx2-Wls cKO cochleae (n, np). Scale bar: 20 µm in (b, c), 200 µm in (d), 5 µm in (g, h), 10 µm in (m-p), 5 µm in (k-k”, l-l”), 2 µm in (k’’’, l’’’). Data are presented as mean ± S.D. and compared using an unpaired t-test. n, number of hair cells measured. Circular mean and variance of hair cell orientation are indicated with blue and red lines in circular histograms, respectively. A Bayesian mixed model analysis used for comparison of circular mean of hair cell orientation between control and cKO groups (see Materials and Methods section for more details). *significantly different (for hair cell orientation analysis), **p < 0.01, ****p < 0.0001, ns = not significant.

### *In silico* analysis identified candidate Wnt ligands from the cochlear epithelium and periotic mesenchyme regulating planar cell polarization

To identify candidate Wnt ligands in the developing cochlea that could contribute to hair cell polarity, we analyzed established single-cell transcriptomic data from the embryonic cochleae. We integrated existing single-cell RNA sequencing data into two categories, each comprised of both epithelial and mesenchymal cells: (1) early (E13.5 and E14.5)^38, 39^ and late embryonic cochlea (E15.5 and E16.5)^39, 40^. From the analysis of both the early and late embryonic cochlea datasets, we identified six cell clusters: hair cells (HCs), cochlear roof, lesser epithelial ridge (LER), greater epithelial ridge (GER), pro-sensory cells, and periotic mesenchyme cells (POM). These clusters were annotated using previously described marker genes^39, 40^.

We next computationally inferred cell-cell communication among different cell clusters with a focus on Wnt ligand-receptor interactions in the developing cochlea using the CellChat^41^ algorithm. First, we combined the cochlear epithelial cells (cochlear roof, LER, and GER cells), while excluding hair cells and prosensory cells. As a result, we generated four new cell groups for the early and late embryonic cochlea datasets: 1) the “Epithelial cells” group (comprising cochlear roof, LER, and GER cells); 2) the “POM” group; 3) the “Prosensory cells” group; and 4) the “Hair cells” group, which includes both IHCs and OHCs (Fig. 2a, b). We then used these newly defined cell groups to create circle plots for each dataset, illustrating significant Wnt-receptor interactions among the cell groups, with the width of the arrows reflecting the scale of these interactions based on the calculated communication probability (Fig. 2c, d)^41^. In the early embryonic cochlea dataset, significant bidirectional Wnt ligand-receptor interactions were detected among all cell groups, along with autocrine signaling (Fig. 2c). In contrast, in the late embryonic cochlea dataset, outgoing Wnts from the hair cell group interacting with any other group were no longer significant (Fig. 2d). Furthermore, in both datasets, the strength of Wnt ligand-receptor interactions involving the POM group was particularly notable (Fig. 2c, d). To identify the Wnt subtype received by the hair cell and prosensory/supporting cell groups, we analyzed the Wnt-ligand interactions in which these cells act as receptors and created chord plots illustrating the significant Wnt ligand-receptor pairs for each dataset (Fig. 2e, f). The values for the relative contribution, reflecting the calculated communication probability of each Wnt ligand-receptor (Source data 1), and are proportional to the width of the lines in the chord plots (Fig. 2e, f). We then categorized and ranked the candidate Wnt ligands received by the hair cell and prosensory/supporting cell groups separately based on whether they originate from the cochlear epithelium or POM (Fig. 2g-j). In the early embryonic cochlea dataset, Wnt7b was identified as the dominant Wnt from the cochlear epithelium, followed by Wnt7a and Wnt5a. In contrast, for the late embryonic cochlea dataset, Wnt5a emerged as the dominant candidate Wnt from the cochlear epithelium, followed by Wnt7b and Wnt7a (Fig. 2g, h). Among the Wnts derived from the POM, Wnt5a was the most highly expressed across both datasets, with Wnt5b following (Fig. 2i, j). In summary, the cell-cell communication analysis of both datasets suggests Wnt5a, Wnt7a, and Wnt7b as candidate Wnts secreted from the cochlear epithelium, with Wnt5a being the most robust among those derived from the POM, that can contribute to Wnt signaling in hair cells during cochlear development.

**Figure 2.**
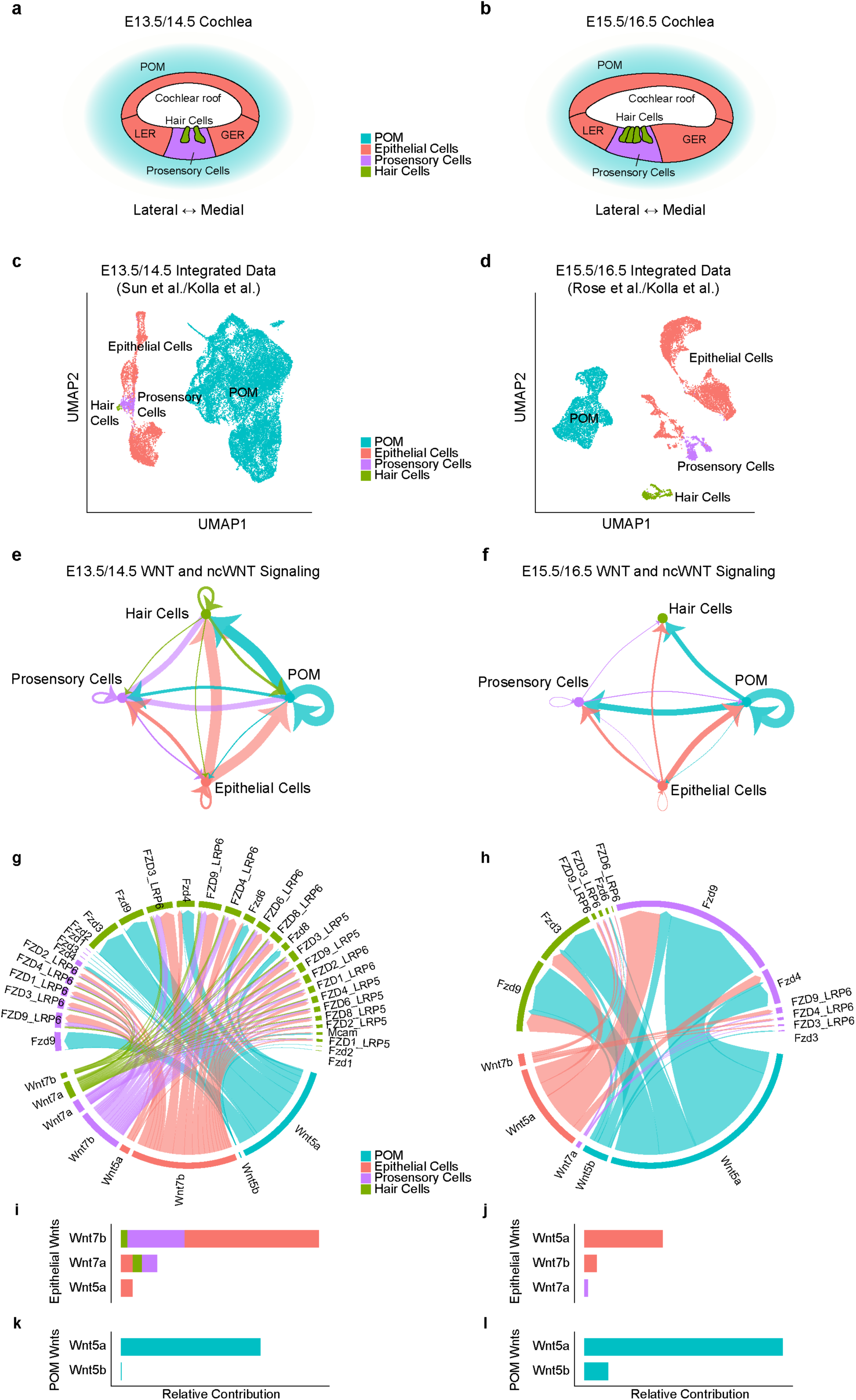
Cell-cell communication analysis of the embryonic cochlea. **a-b.** UMAP of E13.5 periotic mesenchyme (POM) and E14.5 cochlear epithelium (early embryonic cochlea dataset (a) and of E15.5 POM and E16.5 cochlear epithelium (late embryonic cochlea dataset (b). **c-d.** Circle plot showing an inferred signaling network between Wnt (canonical and non-canonical) ligands and receptors among cell groups from the early (c) and late (d) developing cochlea datasets. The width of arrows is proportional to the probability of communication between any two cell groups and autocrine signaling is represented by a loop. In the early dataset, significant Wnt ligand-receptor interactions are found among the four cell groups. Among the significant interactions, outgoing Wnt ligand signals from hair cells are no longer significant in the late embryonic cochlea dataset. Furthermore, the strength of interactions involving POM is robust in both datasets. **e-f.** Chord plots for cell-cell communication from Wnt ligands and receptors in the early (e) and late (f) developing cochlea datasets, with bands’ width reflecting the scale of contribution. **g-j.** Ranked Wnt ligands received by hair cells and prosensory cells in the cochlear epithelium (g, h) or POM (i, j). In the early dataset, Wnt7b is the most potent Wnt ligand emanating from the cochlear epithelium, followed by Wnt7a and Wnt5a (g). In the late dataset, Wnt5a is the most potent Wnt, followed by Wnt7b and Wnt7a (h). Across both datasets, Wnt5a is the most potent among Wnts derived from the POM followed by Wnt5b (i, j). See Source Data file for details.

### Deletion of Wnt5a, Wnt7a, and Wnt7b from the cochlear epithelium shortens cochlea and causes mild hair cell malrotation

Because our computational analysis identified Wnt5a, Wnt7a, and Wnt7b as the candidate Wnt ligands in the cochlear epithelium acting on hair cells and prosensory cells, we hypothesized that concurrent deletion of these Wnt ligands from the cochlear epithelium would phenocopy Emx2-Wls cKO cochlea. To test this hypothesis, we crossed conditional KO animals of Wnt5a^42^, Wnt7a^43^, or Wnt7b^44^ with Emx2-Cre mice, and generated *Emx2^Cre/+^; Wnt5a^flox/flox^* (Emx2-Wnt5a cKO), *Emx2^Cre/+^; Wnt7a^flox/flox^* (Emx2-Wnt7a cKO), and *Emx2^Cre/+^; Wnt7b^flox/flox^* (Emx2-Wnt7b cKO) mice. Previous studies using *in situ* hybridization have demonstrated the expression of Wnt5a, Wnt7a, and Wnt7b in the cochlear epithelium, particularly on the cochlear floor, at E14.5 and E16.5^15, 28, 29^ (Fig. 3a). In E16.5 control cochlea, we confirmed expression of Wnt5a, Wnt7a, and Wnt7b mRNA (Supplementary Fig. 2a, c, e, g, i, k), which were mostly deleted in the cochlear epithelium of the Emx2-Wnt5a, Emx2-Wnt7a, and Emx2-Wnt7b cKO mice (Supplementary Fig. 2b, d, f, h, j, l).

**Figure 3.**
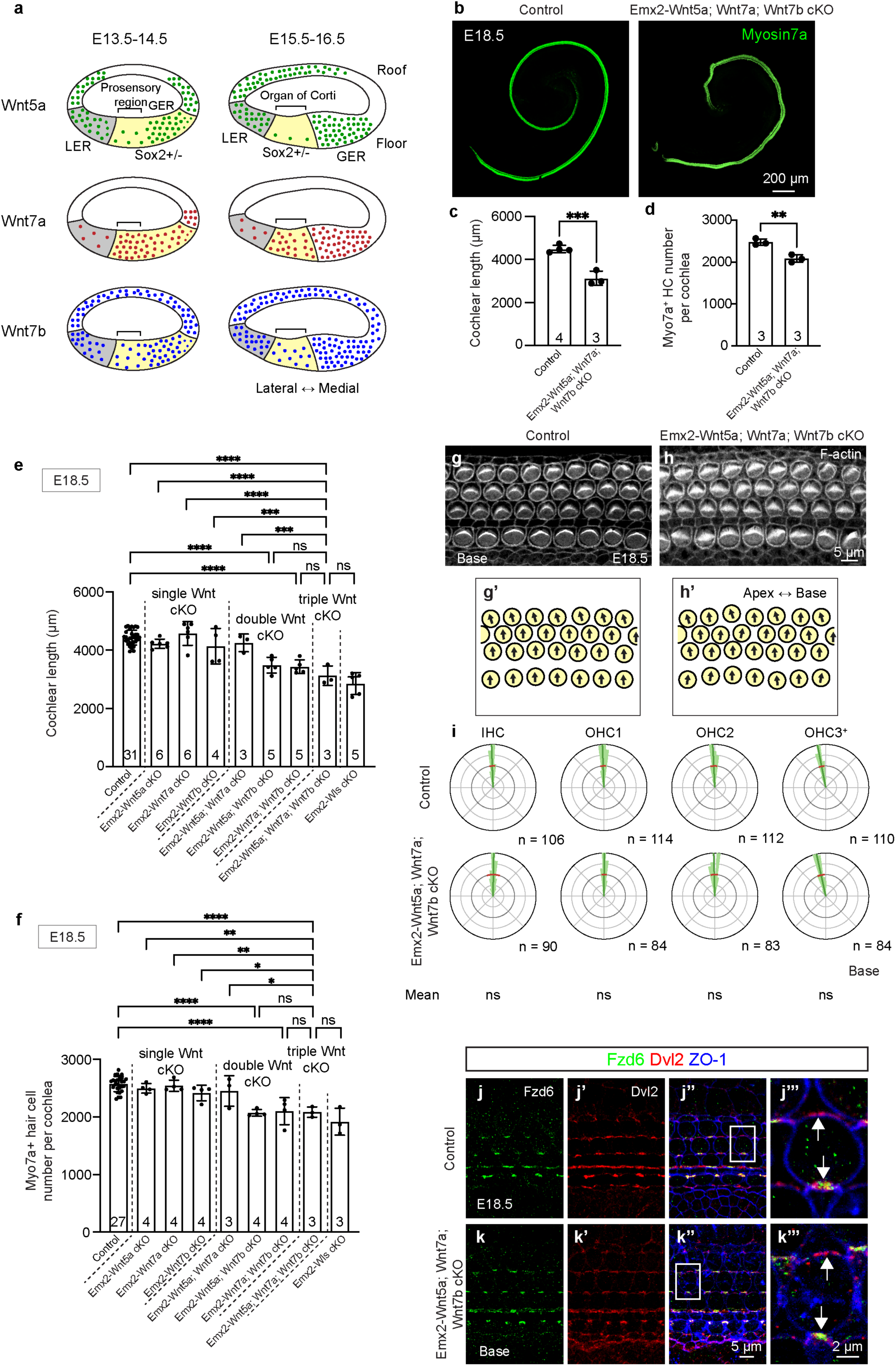
Deletion of multiple Wnts shortens the cochlea and causes mild PCP defects. **a.** Schematic depicting the expression patterns of Wnt5a, Wnt7a, and Wnt7b in the developing mouse cochlear epithelium. **B-C.** Whole mount preparation of E18.5 control and *Emx2^Cre/+^; Wnt5a^flox/flox^; Wnt7a^flox/flox^; Wnt7b^flox/flox^* (Emx2-Wn5a; Wnt7a; Wnt7b cKO) immunostained for Myo7A, with measurements showing that the latter are significantly shorter. **d.** Emx2-Wnt5a; Wnt7a; Wnt7b cKO cochlea had significantly fewer Myo7A-positive hair cells than littermate controls at E18.5. **e**. Measurements of the lengths of cochleae in transgenic mice with deletion of one to multiple Wnts. The length of Emx2-Wn5a; Wnt7a; Wnt7b cKO cochleae was significantly shorter than each of the single or double cKO cochleae, and comparable to that of Emx2-Wls cKO cochleae. **f.** Quantitative analysis of hair cells among control cochleae and cochleae with deletion of one or multiple Wnts. Hair cell counts in Emx2-Wn5a; Wnt7a; Wnt7b cKO cochleae were significantly lower than in each of single or Emx2-Wnt5a; Wnt7a cKO cochleae, and comparable to those in Emx2-Wnt5a; Wnt7b, Emx2-Wnt7a; Wnt7b, or Emx2-Wls cKO cochleae. Hair cell counts in Emx2-Wnt5a; Wnt7b cKO cochleae were significantly lower than Emx2-Wnt5a cKO and Emx2-Wnt7b cKO cochleae (p < 0.001 and p < 0.01, respectively. **g-h.** Whole mount preparation of E18.5 control (g, g’) and Emx2-Wn5a; Wnt7a; Wnt7b cochlea (h, h’) immunostained for f-actin. **i.** Circular histograms illustrating the distribution of hair bundle orientations in control and Emx2-Wn5a; Wnt7a; Wnt7b cKO cochleae. No significant misorientation of the hair cells was detected (circular means and variance). **j–k.** Polarized expression of Fzd6 and Dvl2 is observed at the medial and lateral poles of OHCs, respectively, in the basal turn of both control (j’’’, white arrows) and Emx2-Wn5a; Wnt7a; Wnt7b cKO cochleae (k’’’, white arrows). Scale bar: 200 µm in (b), 5 µm in (g-h, j-j’’, k-k’’), 2 µm in (j’’’, k’’’). Data are presented as mean ± S.D. and analyzed using an unpaired t-test (c, d) or one-way ANOVA followed by either Šídák’s multiple comparisons test (e-f). n, number of hair cells measured. Mean and S.D. of hair cell orientation are indicated with blue and red lines in circular histograms, respectively. A Bayesian mixed model analysis used for comparison of circular mean of hair cell orientation between control and cKO groups. *p < 0.05, **p < 0.01, ***p < 0.001, ****p < 0.0001, ns = not significant.

When Wnt5a, Wnt7a, or Wnt7b was individually ablated in the cochlear epithelium, there was no significant difference in cochlear length compared to littermate controls (p = 0.5132, 0.9070, and 0.4882, respectively; Supplementary Fig. 3a-b, f-g, k-l). Additionally, hair cell counts in these Emx2-Wnt5a and Emx2-Wnt7a cKO cochleae were unchanged (p = 0.5722 and p = 0.5056, respectively), whereas Emx2-Wnt7b cKO cochleae had mildly reduced hair cell counts (∼9%; p < 0.05, Supplementary Fig. 3c, h, m). Furthermore, the circular mean and variance of hair cell orientation in Emx2-Wnt5a and Emx2-Wnt7a cKO cochleae were comparable to controls, while those in Emx2-Wnt7b cKO cochlea were mildly abnormal (significant deviation of IHC, OHC1 and OHC2, Supplementary Fig. 4a-r, Supplementary Table 1). Lastly, Fzd6 and Dvl2 remained polarized in OHCs of Emx2-Wnt5a, Emx2-Wnt7a, and Emx2-Wnt7b cKO cochleae (Supplementary Figure 3d-e’’’, i-j’’’, n-o’’’). These findings suggest that individual deletion of Wnt5a, Wnt7a, or Wnt7b from the cochlear epithelium during development has a minimal effect on cochlear extension and hair cell polarization, corroborating previous reports^15, 30^.

Previously, Wnts were reported to have redundant functions during cochlear development^30, 45^. To rule out this possibility, we generated double KO mice for Wnt ligands: *Emx2^Cre/+^; Wnt5a^flox/flox^; Wnt7a^flox/flox^* (Emx2-Wnt5a; Wnt7a cKO), *Emx2^Cre/+^; Wnt5a^flox/flox^; Wnt7b^flox/flox^* (Emx2-Wnt5a; Wnt7b cKO), and *Emx2^Cre/+^; Wnt7a^flox/flox^; Wnt7b^flox/flox^* (Emx2-Wnt7a; Wnt7b cKO) mice. The cochlear length of Emx2-Wnt5a; Wnt7b and Emx2-Wnt7a; Wnt7b cKO, but not Emx2-Wnt5a; Wnt7a cKO, mice were significantly shorter than those of littermate controls (∼20% and ∼24%, p < 0.01 and p < 0.05, respectively) (Supplementary Fig. 5a-b, f-g, k-l). Hair cell counts in Emx2-Wnt5a; Wnt7b and Emx2-Wnt7a; Wnt7b cKO cochleae were significantly lower than controls (∼20% and ∼15%, p < 0.01 and p < 0.05, respectively), whereas hair cell counts in Emx2-Wnt5a; Wnt7a cKO cochleae were comparable to littermate control cochleae (p = 0.6008, Supplementary Fig. 5c, h, m). Emx2-Wnt5a; Wnt7a cKO cochleae showed normal hair cell orientation (Supplementary Fig. 6a-f, 13), while Emx2-Wnt5a; Wnt7b cKO cochleae exhibited more variable orientation of hair cells compared to controls (circular variance ratio: 1.73, Supplementary Table 1). However, the average orientation of hair cells in Emx2-Wnt5a; Wnt7b cKO cochleae were not significantly different from controls (Supplementary Fig. 6g-r, 13, Supplementary Table 1). Hair cells in all three double cKO cochleae still exhibited polarized expression of Fzd6 and Dvl2 (Supplementary Fig. 5d-e’’’, i-j’’’, n-o’’’), in contrast to the previously reported reduced expression in Emx2-Wnt5a; Wnt7b cKO cochleae^30^.

Finally, we generated and examined the *Emx2^Cre/+^; Wnt5a^flox/flox^; Wnt7a^flox/flox^; Wnt7b^flox/flox^* (Emx2-Wnt5a; Wnt7a; Wnt7b cKO) mice. In these animals, cochlear length was significantly lower than that of littermate controls (∼30%, p < 0.001; Fig. 3b, c), and that after concurrent deletion of Wnt5a and Wnt7a (p < 0.0001, Fig. 3e). Length of Emx2-Wnt5a; Wnt7a; Wnt7b cKO cochlea is comparable to that of Emx2-Wnt5a; Wnt7b cKO, Emx2-Wnt7a; Wnt7b cKO, and Emx2-Wls cKO cochleae (p > 0.05 for all, Fig. 3e). Hair cell counts of Emx2-Wnt5a; Wnt7a; Wnt7b cKO cochleae were significantly reduced compared to littermate controls (∼16% and p < 0.01; Fig. 3d). Similarly, hair cell counts in Emx2-Wnt5a; Wnt7a; Wnt7b cKO cochleae were significantly lower than those in any single cKO (∼17% and p < 0.01 for Wnt5a cKO, ∼18% and p < 0.01 for Wnt7a cKO, and ∼14% and p < 0.05 for Wnt7b cKO cochleae) or Emx2-Wnt5a; Wnt7a cKO cochleae (∼15% p < 0.05), yet comparable to those in Emx2-Wnt5a; Wnt7b cKO, Emx2-Wnt7a; Wnt7b cKO, and Emx2-Wls cKO cochleae at E18.5 (p > 0.05 for all, Fig. 3f). These results suggest that Wnt5a, Wnt7a, and Wnt7b in the cochlear epithelium are individually dispensable yet contribute additively to cochlear outgrowth. By contrast, no misorientation of the hair cells was detected (basal or middle turns) (Fig. 3g-i, Supplementary Fig. 7a-c, Supplementary Table 1). Furthermore, Fzd6 and Dvl2 expression in hair cells remains polarized (Fig. 3j-k’’’). While the length of Emx2-Wnt5a; Wnt7a; Wnt7b cKO is similar to that of Emx2-Wls cKO, hair cell orientation and core PCP protein expression were remarkably normal.

In summary, these data indicate that the effects of Wnt5a, Wnt7a, and Wnt7b from cochlear epithelium are additive for cochlear outgrowth, but not for hair cell orientation. Furthermore, since core PCP protein expression remains polarized in all single, double, and triple conditional KO cochleae, we postulate that additional Wnt ligands, either within or outside the cochlear epithelium, may also contribute to planar polarization.

### Deletion of Wnt5a from the POM does not cause cochlear PCP defects

Our computational cell-cell communication analysis inferred that Wnt ligands from the POM significantly interact with Wnt receptors on hair cells and prosensory/supporting cells, particularly Wnt5a (Fig. 2e, f, i, j). Indeed, a previous study has reported Wnt5a expression in the POM at E14.5^46^. To test the hypothesis that Wnt ligands from the POM regulate cochlear outgrowth and hair cell polarity, we generated *Dermo1 (Twist2)-Cre; Wls^flox/flox^* mice. Dermo1-Cre mice exhibit Cre activity beginning around E9.5 in the mesenchyme-derived tissues^47^, including the POM but not the cochlear epithelium^48^. However, *Dermo1^Cre/+^; Wls^flox/flox^* (Dermo1-Wls cKO) were embryonically lethal (8 of 8 animals by E18.5) as previously reported,^49^ precluding examination of their cochlea. We next generated and examined *Dermo1^Cre/+^; Wnt5a^flox/flox^* (Dermo1-Wnt5a cKO) mice. In control cochleae, Wnt5a mRNA was detected both in the cochlear epithelium and POM (Fig. 4b-b’, Supplementary Fig. 8a-a’, c-c’), corroborating our previous study^15^. By contrast, in Dermo1-Wnt5a cKO cochleae, *Wnt5a* mRNA signals in the POM were undetectable while those in other regions (cochlear epithelium and spiral ganglion) remained present (Fig. 4c-c’, Supplementary Fig. 8b-b’, d-d’), indicating efficient and specific deletion of Wnt5a mRNA from the POM. For the cochlear PCP readouts, however, Dermo1-Wnt5a cKO cochleae exhibited normal length, hair cell counts, hair cell orientation (both circular variance and means), and polarized expression of the core PCP proteins Fzd6 and Dvl2 (Fig. 4d-k’’’, Supplementary Fig. 8e-g, Supplementary Table 1). These results suggest that deletion of Wnt5a alone from the POM does not disrupt cochlear outgrowth, hair cell orientation and PCP signaling.

**Figure 4.**
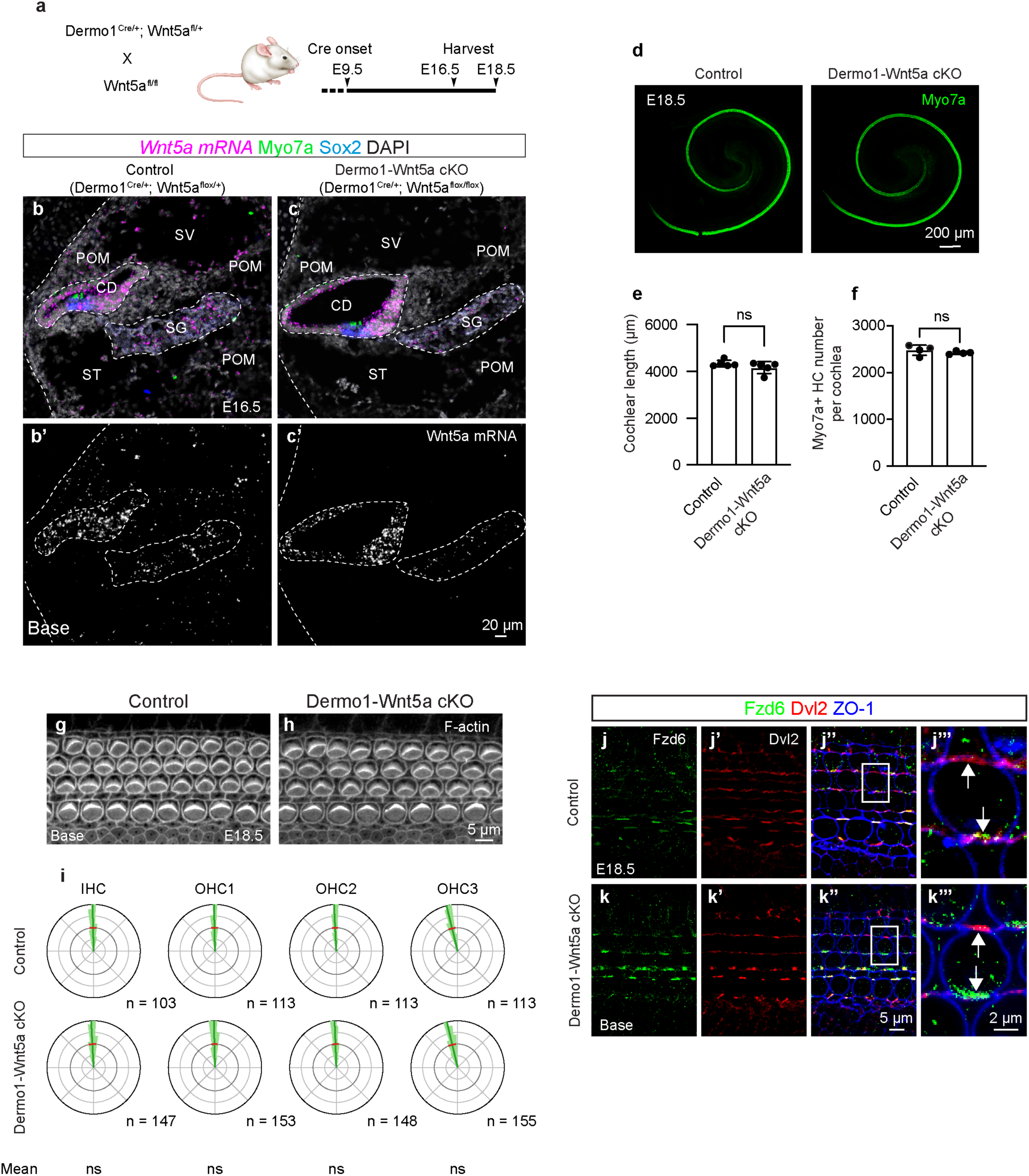
Wnt5a deletion from the periotic mesenchyme does not induce cochlear PCP defects. **a.** Transgenic approach to delete Wnt5a in the POM. Cre recombinase in the POM of Dermo1-Cre cochlea begins around embryonic day (E) 9.5. **d.** Whole mount preparation of cochleae from E18.5 control and Dermo1-Wnt5a cKO embryos. **e-f.** Cochlear lengths and hair cell counts are comparable between control and Dermo1-Wnt5a cKO mice at E18.5. **g-h.** F-actin-stained cochleae from control (g) and Dermo1-Wnt5a cKO (h) embryos at E18.5. **i** Circular histograms demonstrating comparable hair bundle orientation in the basal turn of control and Dermo1-Wnt5a cKO cochleae. **j–j’’’, k–k’’’.** Polarized expression of Fzd6 and Dvl2 (white arrows) is observed at the medial and lateral poles of OHCs, respectively, both in the control (j’’’) and the Dermo1-Wnt5a cKO cochlea (k’’’).These results suggest that deletion of Wnt5a alone from the POM does not disrupt cochlear PCP. CD cochlear duct; ST scala tympani; SV scala vestibule; SG spiral ganglion. Scale bar: 20 µm in (b-c’), 5 µm in (g-h, j-k’’’), 200 µm in (d). n, number of hair cells measured. Mean and S.D. of hair cell orientation are indicated with blue and red lines in circular histograms, respectively. A Bayesian mixed model analysis used for comparison of circular mean of hair cell orientation between control and cKO groups. ns = not significant.

### Inhibition of Wnt secretion in both the cochlear epithelium and periotic mesenchyme causes severe planar polarization defects

When Wnt secretion was prevented in the cochlear epithelium using Emx2-Wls cKO mice, we have shown that hair cell orientation and polarization of two core PCP proteins were disrupted, albeit to a markedly milder degree relative to other PCP mutants^10, 12^ (Fig. 1g-i, Supplementary Fig. 1i-k). Since Wnts from both the cochlear epithelium and POM were predicted to significantly interact with Wnt receptors on hair cells and prosensory/supporting cells (Fig. 2g-q), we generated *Sox9^CreERT^*^2^*; Wls^flox/flox^* (Sox9-Wls cKO) mice to prevent Wnt secretion in both regions. Sox9 is expressed in both the otic/cochlear epithelium and POM from E9.5 to E14.5, before becoming more restricted later^50, 51^. Sox9-Wls cKO animals received tamoxifen (0.2 mg/g of body weight) at E10.5-11.5 or E11.5-12.5 with survival rates of ∼73% at E18.5, consistent with previous reports^52, 53^ (n=48, Fig. 5a, Supplementary Fig. 9a). In E18.5 Sox9-Wls cKO cochlea, we found efficient and broad deletion of Wls in both the cochlear epithelium and POM (Supplementary Fig. 10a-f’).

**Figure 5.**
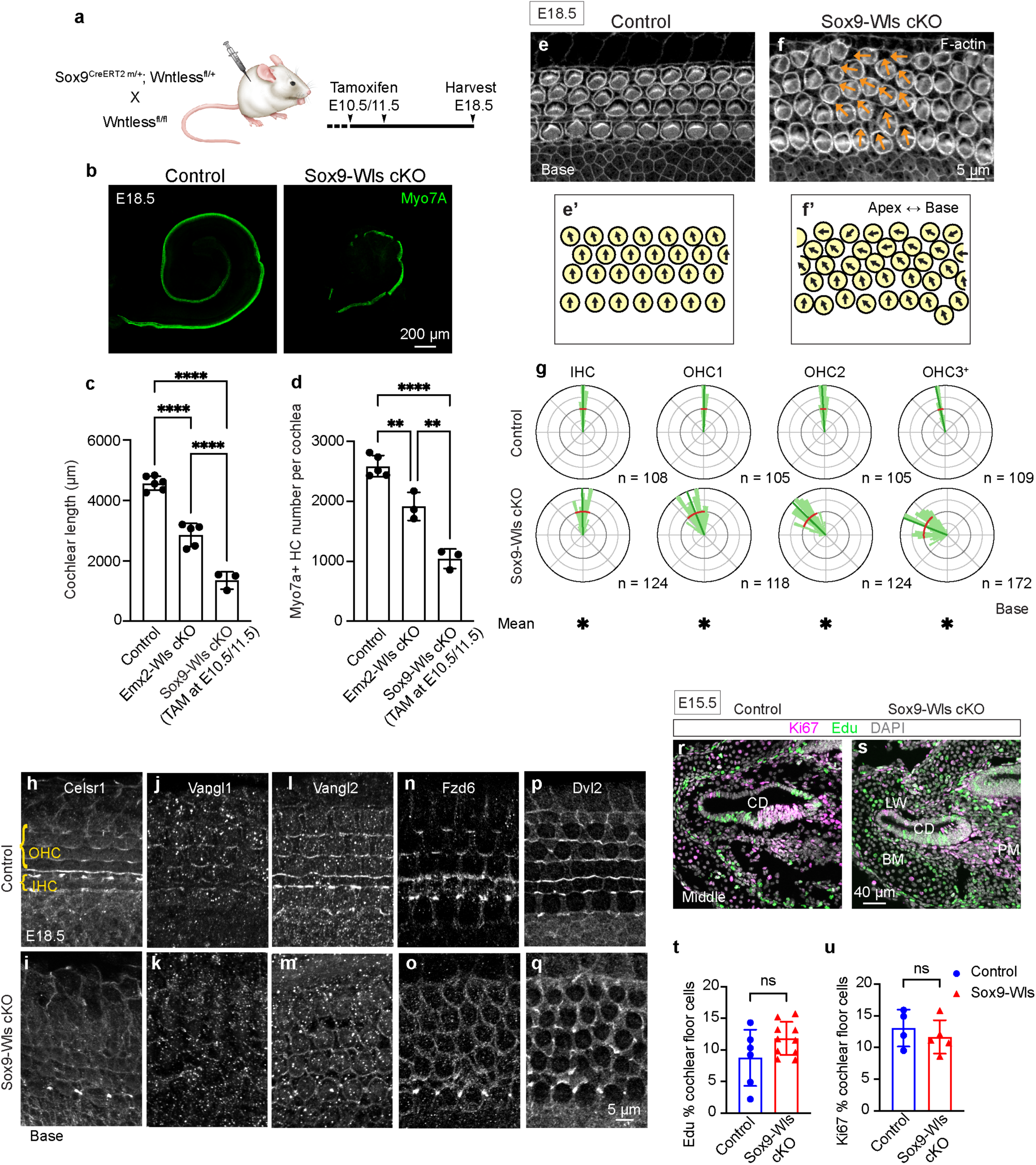
Deletion of Wntless in the cochlear epithelium and periotic mesenchyme causes severe cochlear shortening and PCP defects. **a.** Schematic of deleting Wntless using the Sox9-CreERT2; Wntless-flox mice. Tamoxifen (0.2 mg/g) was administered at E10.5 and E11.5 and cochleae were analyzed at E18.5. **b-c.** Immunostaining and measurements of E18.5 control (*Sox9^CreERT^*^2^*^/+^; Wls^flox/+^*) and *Sox9^CreERT^*^2^*^/+^; Wls^flox/flox^* (Sox9-Wls cKO) cochleae, showing dramatic shortening of the latter and significantly shorter than both control and Emx2-Wls cKO cochleae. **d.** Sox9-Wls cKO cochleae had significantly fewer cells compared to control and Emx2-Wls cKO cochleae. **e-f.** Whole mount preparation of basal turn of the E18.5 f-actin-stained cochlea The Sox9-Wls cKO cochlea shows five to six rows of hair cells that are disorganized and lack a separation between IHCs and OHCs. Notably, hair cells are mostly malrotated toward the apex. **g** Circular histograms illustrating the distribution of hair bundle orientation in control and Sox9-Wls cKO cochleae, which showed significantly larger circular variance and different average orientation in each row relative to controls. **h-q.** Immunostaining for core PCP proteins in the control and Sox9-Wls cKO cochleae for Celsr1 (h, i), Vangl1 (j, k), Vangl2 (l, m), Fzd6 (n, o), and Dvl2 (p, q), showing a loss of polarized expression of Celsr1, Vangl1, Vangl2, Fzd6, and Dvl2 in OHCs of Sox9-Wls cKO cochleae. **r-s.** Sections of E15.5 control (r) and Sox9-Wls cKO (s) cochleae illustrating Ki67- and EdU-positive cells in both the cochlear epithelium and periotic mesenchyme. **t-u.** Quantitative assessment showing comparable proliferation in the cochlear epithelium between the control and Sox9-Wls cKO cochleae. CD, cochlear duct; LW, POM near the lateral wall; BM, POM near the basilar membrane; PM, perimodiolar POM. Scale bar: 200 µm in (b), 5 µm in (e-f, h-q), 40 µm in (r-s). Data are presented as mean ± S.D. and analyzed using one-way ANOVA followed by Tukey’s multiple comparisons test (c-d) and an unpaired t-test (t-u). n, number of hair cells measured. Mean and S.D. of hair cell orientation are indicated with blue and red lines in circular histograms, respectively. A Bayesian mixed model analysis used for comparison of circular mean of hair cell orientation between control and cKO groups. *significantly different (for hair cell orientation analysis), **p < 0.01, ****p < 0.0001, ns = not significant.

After tamoxifen administration at E10.5-11.5, Sox9-Wls cKO cochleae were dramatically shorter and contained significantly fewer hair cells than controls (∼70% and ∼60%, p < 0.0001 for both, Fig. 5b-d) and Emx2-Wls cKO cochleae (∼38% and p < 0.0001, and ∼46% and p < 0.01, respectively). Sox9-Wls cKO cochlea displayed five to six rows of hair cells throughout the cochlea (Fig. 5e-f, Supplementary Fig. 11a-b), consistent with a defect in convergent extension seen in PCP mutants^54, 55^. Additionally, hair cells appeared disorganized without a clear separation between IHCs and OHCs (Fig. 5f, Supplementary Fig. 11b), possibly suggesting abnormal pillar cell development. Furthermore, hair cells were severely malrotated with significantly higher variance and more deviation than controls (circular variance ratio: 5.33, Fig. 5e-g, Supplementary Fig. 11a-c, Supplementary Table 1). Unlike Emx2-Wls cochleae, where only a subset of hair cells was significantly malrotated and directed to the base, Sox9-Wls cochleae shows that each row of hair cells is severely malrotated (Fig. 1g-i, 4e-g, Supplementary Fig. 1i-k, 11a-c, Supplementary Table 1). In contrast to PCP mutants where hair cells are randomly malrotated^12, 16, 17^, Sox9-Wls cochlear hair cells are rotated towards the apex, suggesting a global instructive role for secreted Wnts.

In control cochleae, Celsr1, Vangl1, Vangl2, and Fzd6 are localized to the medial junctions of hair cells, while Dvl2 is found at the lateral junctions (Fig. 5h, j, l, n, p). In contrast, each of these core PCP proteins is mislocalized in hair cells in Sox9-Wls cKO cochleae (Fig. 5i, k, m, o, p). Overall, compared to Emx2-Wls cKO cochleae, Sox9-Wls cKO mice exhibited greater disruptions in cochlear duct elongation, hair cell orientation, and polarization of core PCP proteins in hair cells.

To determine whether the reduced length and hair cell counts in Sox9-Wls cKO cochleae were attributed to reduced proliferation, we performed Ki67 immunostaining and EdU labeling. EdU (50 mg/kg, IP) was administered at E12.5, along with tamoxifen (0.2 mg/g) at E10.5-11.5. At E15.5, both control and Sox9-Wls cKO cochleae displayed Ki67-positive and EdU-positive cells in both the cochlear epithelium and POM (Fig. 5r-s). No significant difference was observed in the number of EdU- or Ki67-labeled Sox2-positive cells in control and Sox9-Wls cKO cochleae (p = 0.1015 and p = 0.4776, respectively; Fig. 5t-u, Supplementary Fig. 11d-e). Additionally, no significant differences were detected in the number of EdU- or Ki67-positive cells in the POM (p = 0.7144 and p = 0.4975, respectively; Fig. 5r-s, Supplementary Fig. 11d-e). These findings suggest that the cochlear shortening and reduced cochlear hair cell counts observed in Sox9-Wls cKO mice are primarily due to disruptions in cochlear outgrowth and/or convergent extension.

To determine whether the regulation of cochlear outgrowth and PCP development by Wnt secretion is time-dependent during cochlear development, we ablated Wntless one day later (tamoxifen at E11.5-12.5, Supplementary Fig. 9a). Sox9-Wls cKO cochleae were still significantly shorter (∼26% and p < 0.05) and contained fewer hair cells (∼14% and p < 0.05) compared to controls at E18.5 (Supplementary Fig. 9b-d). However, when compared an earlier injection of tamoxifen at E10.5-11.5, both the cochlear length (∼60% and p < 0.0001) and hair cell counts (∼53% and p < 0.0001) were significantly higher (Supplementary Fig. 9c-d). Additionally, hair cells remain significantly malrotated towards the apex and showed a high variance in orientation (circular variance ratio: 3.28, Supplementary Fig. 9e-j, Supplementary Table 1). However, core PCP proteins (Fzd6, Celsr1, Vangl1, and Vangl2) were all polarized in hair cells (Supplementary Fig. 9k-s), suggesting the overall effects were less severe than Wntless deletion earlier. These results suggest that the effect of Wnts on the cochlear PCP development is highly time-sensitive during the embryonic period.

Overall, these data suggest that secreted Wnt proteins, around E10.5-11.5, from both the POM and cochlear epithelium are required for elongation of the cochlear duct, orientation of hair cells, and polarization of core PCP proteins in hair cells during cochlear development.

## Discussion

During development, the mouse cochlea elongates to form a spiral organ that is two and a half turns in length, in which sensory and non-sensory cells differentiate and undergo precise radial patterning. These steps are temporally and spatially coordinated and governed by the Notch, Wnt, and Shh signaling pathways^2^. PCP signaling critically regulates cochlear elongation and sensory cell polarization. Previous studies have shown that secreted Wnts in the cochlear epithelium are required for cochlear elongation, sensory cell polarization, and polarization of PCP proteins^15, 31^, although the severity of the sensory cell polarization defects was remarkably mild, raising the possibility that other factors co-regulate these steps. Here, based on computational analysis of single-cell transcriptomes of the developing cochlear epithelium and surrounding POM, we predicted that Wnt5a, Wnt7a and Wnt7b in the cochlear epithelium, and Wnt5a in the POM were the most potent Wnts acting on the developing hair cells and prosensory/supporting cells. Using combinations of transgenic mice, we selectively deleted Wnt members or Wnt secretion in a spatiotemporally selective manner. We first demonstrated that these Wnt members in both the cochlear epithelium and POM additively contribute to cochlear outgrowth and convergent extension. By contrast, hair cell orientation and PCP signaling based on the polarized expression of core PCP proteins are only mildly affected despite deletion of all three Wnt members or Wntless in the cochlear epithelium. When Wnt secretion was prevented in both the cochlear epithelium and POM, cochlear length, hair cell orientation and PCP signaling were severely affected, suggesting that Wnt members distributed across the epithelial and mesenchymal compartments are highly redundant in regulating cochlear development.

### Wnts are required for cochlear hair cell planar cell polarity and cochlear outgrowth

In our current study, inhibition of Wnt secretion in the cochlear epithelium during development only caused mild hair cell misorientation and loss of polarization in a subset of core PCP proteins, consistent with previous studies^15, 31^. This suggests that Wnt proteins from the cochlear epithelium have a limited role in establishing hair cell polarity. In contrast, Sox9-Wls cKO cochleae demonstrated severe hair cell misorientation and loss of polarization of all core PCP proteins examined. Thus, Wnts from both the cochlear epithelium and POM are critical in regulating hair cell polarity.

Wnts have previously been implicated to function as global cues for PCP^25^. In Drosophila, Wingless and Wnt4 modulate intercellular Fzd–Vangl interactions, thereby establishing directional cues for PCP axis orientation^56^. However, recent studies in Drosophila have challenged the idea that secreted Wnts are required for core PCP protein enrichment and planar polarity^57, 58^. In various vertebrate organs, the role of Wnt ligands in regulating PCP is well established. For instance, Wnt5b and Wnt11 coordinate cardiac remodeling through polarization of actomyosin activity in zebrafish. During chick muscle development, Wnt11 directs elongation of myocytes along the antero-posterior axis of the embryo^59^. In mice, Wnt5a establishes PCP in limb mesenchyme^60, 61^ and Wnt5a and Wnt5b direct node cell polarization in the developing heart^62^, while Wnt5a induces phosphorylation of Vangl2 in a dose-dependent manner^63^. However, the relationship between Wnt signaling and PCP is not yet clear in the cochlea.

In the mouse cochlea, several studies have found that Wnt proteins in the cochlear duct regulate the orientation and PCP signaling in hair cells^15, 30, 31^. However, in part because deficiency of Wnt proteins results in hair cell defects that are relatively mild compared to other PCP mutants, it has been suggested that Wnts do not play a central role in determining cochlear PCP^32, 33^. Our results in this study clearly show that both epithelial and mesenchymal Wnts contribute to cochlear outgrowth and hair cell planar cell polarity, and not merely epithelial Wnts as previously suggested^15, 30, 31^.

PCP mutants such as *Vangl2^Lp/Lp^* demonstrate convergent extension defects, with shorter cochleae (∼27% of wild-type littermates) without a significant reduction of hair cells^10^. In contrast, ablation of Wntless in the cochlear epithelium (Emx2-Wls cKO) and in both the cochlear epithelium and POM (Sox9-Wls cKO) lead to significantly shorter cochleae (∼38% and 70%, respectively) and a decrease in hair cells. Because the degree of shortening is severe and more than accounted for by the convergent extension defect alone, our study suggests that both epithelial and POM Wnt proteins are required for cochlear outgrowth and convergent extension. When comparing the results from transgenic mouse lines to delete one or multiple Wnts and epithelial or both epithelial and mesenchymal Wnts, we conclude that multiple Wnt members additively contribute to cochlear outgrowth (Fig. 6).

**Figure 6.**
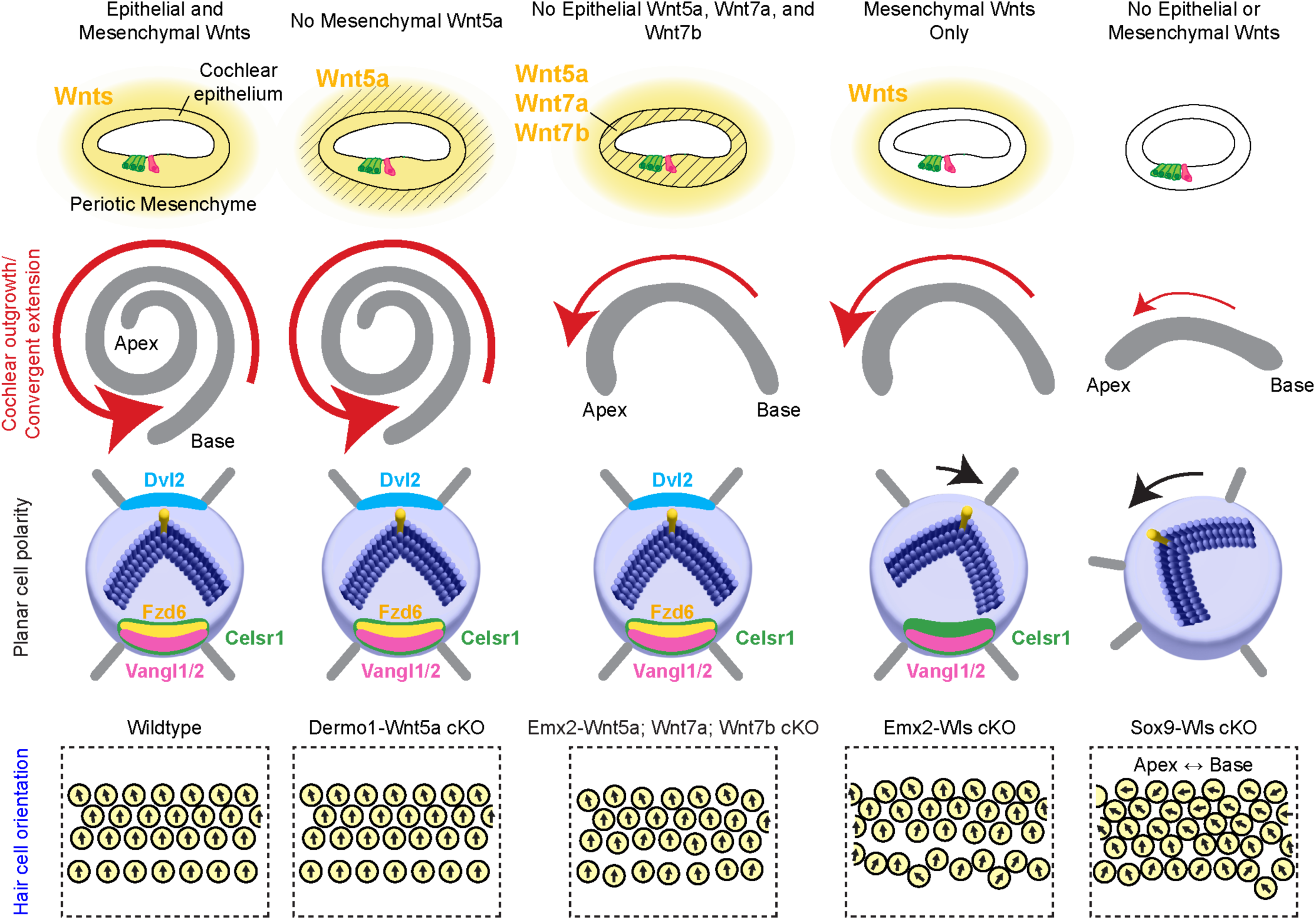
Working model of epithelial and mesenchymal Wnts coordinating to regulate cochlear elongation, hair cell orientation, and PCP signaling. When only mesenchymal Wnts are present, cochlear length is mildly shortened and hair cells mildly misoriented and most core PCP proteins are present. In the absence of both epithelial and mesenchymal Wnts, cochlear duct is severely shortened, and hair cells severely malrotated toward the apex.

While cochlear PCP mutants are characterized by hair cell malrotation, different degrees of defects have been reported, with Vangl2 and Celsr1 mutants being more severe and the Vangl1 mutant milder^16, 17^. Moreover, core PCP proteins serve redundant functions, with deletion of multiple members or the presence of the dominant negative Vangl2-looptail mutation accentuating the polarity defects^12, 14, 16^. Several studies have demonstrated that deletion of Wntless or multiple Wnts in the cochlear epithelium causes mild hair cell polarity defects and loss of or decreased polarized expression of Fzd6 and Dvl2, but not other core PCP proteins. When Wnt secretion is prevented in both cochlear epithelium and periotic mesenchyme (using Sox9-Wls cKO at E10.5-11.5 and not E11.5-12.5), hair cell polarity defects are severe and all core PCP proteins mislocalized. Thus, our study demonstrates that secreted Wnt proteins are not only required for PCP establishment in the developing cochlea (hair cell orientation and core PCP localization) but also do so in an additive and time-dependent manner.

### Dual regulation of hair cell orientation by PCP core proteins and Wnts

Cochlear hair cells are uniformly oriented with their bundles facing laterally. Prior work on PCP mutants has characterized hair cell orientation to be highly variable with random rotation in both clockwise and counterclockwise directions^10, 12, 17^. These results suggest a local, instructive role for core PCP proteins in guiding the polarization of hair cells and limiting variance in orientation. In support of this role, Vangl2 misexpression increases the variance of hair cell orientation without affecting the average angle of their rotation in the chicken basilar papilla^64^.

The role of Wnt ligands in regulating hair cell orientation and PCP is less clear, as Wntless deletion in the cochlear epithelium increases variance in hair cell orientation without significantly affecting the average position, suggesting also a local instructive role^15^. However, in the presence of the Vangl2-looptail mutation, Wntless deletion in the cochlear epithelium leads to malrotation of hair cells towards the apex, suggesting that Wnt ligands in the epithelium may serve as an instructive global cue^15^. In a similar vein, inhibition of endogenous Wnts by inhibitors Sfrp1 or Wif1 in cultured mouse cochlea directed hair cells to uniformly rotate unidirectionally^28^. In the current study, deletion of Wls from both the cochlear epithelium and POM resulted in pronounced and uniform hair cell rotation toward the apex, further supporting the notion that Wnt proteins provide an instructive global cue for hair cell orientation. Together, we propose a model where hair cell orientation and cochlear outgrowth are regulated by both core PCP proteins and Wnt proteins from the epithelial and mesenchymal compartments in the developing cochlea. In the model, core PCP proteins serve local instructive roles while Wnt proteins play a more global role in guiding cell and tissue organization (Fig. 6).

In *Vangl2^Lp/Lp^* and other PCP mutants, hair cell malrotation is most severe in the lateral-most most hair cells (i.e., abneural OHC3+), but the direction of malrotation is random^10^. After broad Wntless deletion (Sox9-Wls cKO), hair cell malrotation is also most severe in the lateral-most hair cells (Supplementary Table 1), but malrotation is almost exclusively directed towards the apex. Hair cell malrotation towards the apex is also observed in Emx2-Wls cKO in the presence of the *Vangl2^Lp/+^* allele^15^, suggesting that Wnts may act as a global instructive cue for hair cell orientation. It is possible that other signaling pathways also contribute to cochlear patterning. For example, Wnt and FGF signaling are both crucial in maintaining migratory polarity and gene expression domains in the lateral line primordium in zebrafish^65^ and regulating PCP during mouse limb development^61^. Lastly, it is also possible that local shearing forces organize and pattern hair cells and supporting cells differentially along the abneural-neural axis^32, 66^.

### Wnt ligand-receptor interaction between epithelial and mesenchymal tissues

Our study suggests that Wnt proteins from both the epithelium and POM regulate the development of PCP in the cochlear epithelium. While Wnt ligands are classically thought to act at short to medium ranges, they have been shown to travel remarkable distances to mediate mesenchymal-epithelial crosstalk in other organ systems. For example, Wnt9b in the Wolffian duct epithelium also regulates the development of metanephric mesenchyme and mesenchymal to epithelial transitions^67^. Also, mesenchymal cells secrete Wnt proteins to regulate stem cell proliferation and differentiation in both the intestines and airway^68, 69, 70^. These studies illustrate the Wnt-mediated mesenchymal-epithelial crosstalk during tissue development and homeostasis. The distance and mechanisms by which Wnts are transported are likely context-dependent^71, 72, 73^, and warrants investigation given their significant roles in cochlear development.

### Many Wnt members contribute to cochlear development

Among the seven Wnt members expressed in the developing cochlea, our computational cell-cell communication analysis predicts Wnt5a, Wnt7a, Wnt7b as the most potent ligands to interact with corresponding receptors on hair cells. We and others have found robust expression of Wnt5a mRNA in the cochlear epithelium flanking the organ of Corti and the periotic mesenchyme from E12.5 to E16.5, higher in the base than the apex^15, 30^. On the other hand, Wnt7a and Wnt7b mRNA are both broadly expressed in the cochlear duct with no apparent apical-basal gradient. In agreement with prior studies^15, 30^, deletion of both Wnt5a and Wnt7b in the cochlear duct leads to significant shortening of the cochlear duct and mild hair cell orientation defects. However, either deletion alone has a minimal effect. Interestingly, deletion of both Wnt7a and Wnt7b in the cochlear duct shortens the cochlea to the extent similar to that of deletion of Wnt5a/Wnt7b, supporting the notion that Wnt proteins additively regulate cochlear outgrowth. As Emx2-Wls cKO cochleae display more severe PCP defects than Emx2-Wnt5a; Wnt7a; Wnt7b cochleae, we attribute this to either Wnt redundancy or incomplete deletion in the latter model. Four other Wnt members (Wnt4, Wnt9a, Wnt11, and Wnt2b) are also expressed at low levels in the developing cochlear duct^15^ and may also contribute to hair cell PCP signaling. According to our cell-cell communication computational analysis, Wnt5a is deemed the most potent Wnt ligand; however, neither deletion of Wnt5a in the POM (DermoCre-Wnt5a cKO) nor in the cochlear duct alone^15^ causes cochlear shortening or hair cell misorientation.

### Wnt signaling is not required for hair cell formation

Our data demonstrate that even though we deleted Wnts or Wls either in the cochlear epithelium or in b oth the cochlear epithelium and POM, hair cell formation still occurs. This data suggests that secreted Wnts are not required for hair cell formation after E10.5. Previous *in vivo* mouse studies deleting beta-catenin, the central mediator of canonical Wnt signaling, show conflicting results. In mice where conditional deletion of beta-catenin was performed using the CMV-CreER allele (tamoxifen at E11.5), Atoh1+ hair cells were absent at E14.5^21^. However, hair cells were observed at later time points (E15.5-18.5) when other mouse lines were examined (Sox2-CreERT2 (tamoxifen at E12 or E13), Fgf20-Cre, Emx2-Cre)^21, 23, 74^. Together with the current study, these results suggest that neither beta-catenin nor secreted Wnts are required for cochlear hair cell formation.

### Summary

In conclusion, secreted Wnts from the cochlear epithelium and POM are essential for establishing planar polarity in cochlear hair cells and for cochlear outgrowth during development. Additionally, multiple Wnt ligands, including Wnt5a, Wnt7a, and Wnt7b, contribute to cochlear outgrowth, hair cell orientation and PCP protein localization, to ensure precise cochlear length and hair cell patterning.

## Materials and methods

### Mice

The following mouse strains were used; *Wls-flox*^75^(#012888, Jackson laboratory, Bar Harbor, ME), *Wnt5a-flox*^42^ (Strain #026626), *Wnt7b-flox*^44^(#008467), Sox9-CreERT2^76^ (#018829) (All from the Jackson Laboratory), *Wnt7a-flox*^43^, Emx2-Cre^36^, and *Dermo1(Twist2)-Cre* mice ^77^. All the floxed mouse lines for the Wnts above mentioned are maintained by homozygous sibling matings, while *Emx2^Cre/+^* and *Dermo1^Cre/+^* mouse lines are maintained by breeding heterozygous mice. Tamoxifen was administered intraperitoneally to dams carrying the Sox9-CreERT2 allele at a dose of 200 µg/g. For the EdU assay, a single dose (50 mg/kg; Invitrogen) was given via intraperitoneal injection to pregnant mice at E12.5, and then cochlear tissue was collected at E15.5. Embryos were considered E0.5 at noon on the day when a plug was discovered. The protocol is approved by the Animal Care and Use Committee of Stanford University School of Medicine (#18606).

### Genotyping

DNA templates were generated by incubating tissues (ear punch or tail clipping) in 180 μl of 50 mM NaOH at 98°C for 1 h followed by the addition of 20 µl of 1 M TrisHCl (pH 8.0). We used either GoTaq® Master Mixes (M712) or KAPA2G Fast HotStart® ReadyMix™(KK5609) to amplify DNA fragments. The primers used were: Wls-flox (fwd, 5′-aggcttcgaacgtaactgacc-3′; rev, 5′-ctcagaactcccttcttgaagc-3′), Wnt5a-flox (fwd, 5′-ggtgagggactggaagttgc-3′; rev, 5′-ggagcagatgtttattgccttc-3′), Wnt7a-flox (fwd, 5′-cacagccacccctagagagctcaatt-3′; rev, 5′-atgctttgccaggga acaccc-3′), Wnt7b-flox (fwd1, 5′-gccagaggccacttgtgtag-3′; fwd2, 5′-ggtagtccttcctgcccttt-3′; rev, 5′-gtgtgtcctggcctgatttt-3′), Emx2-Cre and Sox9-CreERT2 (fwd, 5′-gagtaatagcgaccaatcatcaagcc-3′; rev1, 5′-cgaacatcttcaggttctgcgg-3′; rev2, 5′-cttggaagcgatgacccagatatcgg-3′), and Dermo1-Cre (wt fwd, 5′-cgtctcagctacgccttctc-3′; wt rev, 5′-tcacgaggagagatacactgg-3′; mt fwd 5′-aacttcctctcccggagacc-3′; mt rev, 5′-ccggttattcaacttgcacc-3′).

### Immunohistochemistry

The heads of E15.5 or E18.5 mouse embryos were halved and fixed with 4% paraformaldehyde (PFA, Electron Microscopy Sciences) in phosphate-buffered saline (PBS, pH 7.4) for 2 hours for the following markers: Myosin7a, Sox2, Celsr1, Phalloidin, Ki67, and DAPI. Fixation with 10% Trichloroacetic acid (TCA, Sigma-Aldrich, 1 hour on ice) in Milli-Q water was performed for these markers: Frizzled-6 (Fzd6), Dishevelled-2 (Dvl2), Vangl1, Vangl2, and ZO-1. After fixation, all tissues were stored in PBS at 4°C.

For whole-mount preparation of E15.5 and E18.5 mouse cochlea, the otic capsule was dissected from the fixed tissues, followed by the removal of the lateral wall. For cochlear sections, fixed otic capsule was treated with 10%, 20%, and 30% sucrose in PBS at 4°C for 1 hour, 1 hour, and overnight, respectively. The samples were then embedded in OCT for cryosectioning at a thickness of 10 µm. Both the whole-mount preps and sections were incubated with 0.1% Triton X-100 in PBS for 1 hour at room temperature. Following this, the samples were blocked with 5% donkey serum, 0.1% Triton X-100, 1% bovine serum albumin (BSA, #BP1600–100, Thermo Fisher Scientific), and 0.02% sodium azide (NaN3, #S2002–25G, Sigma-Aldrich,) in PBS (1 hour at RT). Tissues were then incubated with primary antibodies overnight at 4°C. The primary antibodies used were: Myosin7a (rabbit, 1:1000, 25-6790, Proteus Bioscience), Fzd6 (goat, 1:250, AF1526, R&D Systems), Dvl2 (rabbit, 1:250, 12037-1-AP, Proteintech, Rosemont, IL), ZO-1 (mouse IgG1, 1:1000, 33-9100, Thermo Fisher Scientific), Ki67 (rat, 1:250, rat, 14-5698-82, Thermo Fisher Scientific), Vangl1 (rabbit, 1:1,000, HPA025235, Sigma-Aldrich), Vangl2 (rabbit, 1:250, 21492-1-AP, Proteintech), and Celsr1 (rabbit, 1:3,000, gift from E. Fuchs, Rockefeller University). Following incubation with primary antibodies, samples were washed with PBS three times (15 minutes each) and incubated with the following secondary antibodies: Alexa Fluor 488, 546, and 647 (1:400 for all, A-11055, A-10040, A-31571, Thermo Fisher Scientific) and DAPI (1:10,000, Invitrogen) at room temperature for 2 hours. To label f-actin, phalloidin conjugated with Alexa546 (1:500, A-22283, Thermo Fisher Scientific) was applied during secondary antibody incubation. EdU detection was performed using the 647 Click-IT kit (C10340, Invitrogen). After staining, cochlear epithelium was mounted in ProLong™ Gold Antifade Mountant (P36930, Thermo Fisher Scientific) and coverslipped. Control and experimental samples were always processed and imaged in parallel.

### Fluorescent RNA *in situ* hybridization

The whole heads of E16.5 or E18.5 mouse embryos were halved and fixed with 4% PFA in PBS for 16 to 24 hours at 4°C, treated with 10, 20, and 30% sucrose in PBS at 4°C (1 hour, 1 hour, and overnight, respectively). After that, tissues were embedded in OCT for cryosections (10 µm). RNAScope and BaseScope assays were performed as follows: tissue sections were hybridized with commercial probes from Advanced Cell Diagnostics (ACDbio). These probes were specifically designed to detect deleted regions of the following genes in that were deleted: Wls, Wnt5a, Wnt7a, and Wnt7b. The sections were counterstained with anti-Myosin7a (rabbit, 1:1000, 25-6790, Proteus Bioscience) and Sox2 (goat, 1:200, AF2018, R&D Systems) antibodies. The use of these antibodies was done according to the manufacturer’s instructions for fixed frozen sections. DAPI (1:10,000, Invitrogen) was also used for nuclear staining. Briefly, sections were washed in PBS for 5 mins once and then treated with H_2_O_2_ for 10 minutes. Next, the sections were permeabilized using a target retrieval reagent (ACDbio) and proteinase prior to hybridization. The following probes were used: DapB (Cat No. 310043), Polr2a (Cat No. 310451), Wls (Cat No. 405011), Wnt5a-2EJ (Cat No. 717251), Wnt7a-3zz-st-C1 (Cat No. 1277921-C1), and Wnt7b-2zz-st-C1 (Cat No. 1275011-C1) (all from Advanced Cell Diagnostics). Control and cKO samples were processed and imaged in parallel.

### Imaging and quantification

Immunofluorescence or *in situ* hybridization samples were imaged using a Zeiss LSM700 or LSM880 confocal microscope in conjunction with Zen software (Zeiss). Image analysis, including measurements of cochlear length and hair cell orientation, was performed using ImageJ software as described below (NIH).

Wholemount preparation and cochlear section were used to assess proliferation in the cochlear epithelium and of periotic mesenchymal cells, respectively. For the former, Sox2+ cells in the cochlear epithelium in the basal, middle, and apical regions were used to calculate the percent Edu+ or Ki67+. For the POM, 20-50 DAPI+ cells from three areas (lateral wall, perimodiolar, and basilar membrane) in the basal, middle, and apical regions of the cochlea were used to calculate of the percent of Edu+ or Ki67+ cells. The basal, middle, and apical regions were defined as 25, 50, and 70% length from the basal end of the cochlear epithelium.

### Hair cell orientation measurement

Whole mount preparation of the basal and middle turns of the right cochlea was used. The orientation of individual hair cells was defined as previously described^15^. Specifically, using ImageJ, a line was drawn parallel to the pillar cells (a reference line), and another line was drawn from the base to the apex of bundles of individual hair cells (a hair bundle line). Then, the angle between the hair bundle and reference lines was used to determine the orientation of the hair cell (in degrees, counterclockwise direction is positive). Thirty to 50 hair cells for each row from one cochlea from each mouse were measured (IHC, OHC1, OHC2, OHC3, and additional rows more lateral to the OHC3 are denoted as OHC3+). Three to six mice for each genotype were examined. Polar coordinate histogram plots depicting hair cell orientation at each row and condition were generated and statistics including the circular mean, mean resultant length, standard deviation, and variance were calculated in using Python and the code is available on GitHub (https://github.com/tahajanlab/published/tree/main/Kishimoto_et_al).

### Inferred Cell-Cell Communication Analysis

To include both epithelial cells and mesenchymal cells in the cell-cell communication analyses, we merged previously published E14.5/E16.5 epithelial-enriched data (GEO Accession GSE137299^39^) with E13.5/E15.5 mesenchymal-enriched datasets (GEO Accession GSE178931^38^; GEO Accession GSE217727^40^) in Seurat v5.3.0 merge function resulting in combined raw count matrices^78^. The merged data was then log transformed, normalized, and scaled for sequencing depth. Linear dimensionality reduction was then performed via principal component analysis (PCA). For visualization, integration was performed using Harmony^79^ and non-linear dimensionality reduction was performed via the uniform manifold approximation and projection (UMAP) algorithm. Cell types were identified by canonical markers for mesenchymal cells (*Pou3f4*), prosensory cells (*Sox2* and *Cdkn1b*), hair cells (*Pou4f3*), and the rest of the epithelial cells (*Epcam*). CellChat was then used to infer cell-cell communication probability based on ligand and receptor gene expression. Normalized count data was imported into CellChat v2.1.2 and the CellChatDB mouse database was used^41^. Communication probability was calculated based on the 10% truncated mean (average expression after discarding 10% from each end of the data) for each ligand and receptor gene. The p-values were Benjamini-Hochberg adjusted, and a cutoff of an adjusted p-value < 0.05 was considered statistically significant. Chord plots were used to visualize directed ligand-receptor interactions for the WNT and non-canonical WNT signaling pathways with a specific focus on hair cells and prosensory cells as signal receivers. Bar graphs were also used to show relative contributions of cell signaling from hair cells, prosensory cells, and epithelial cells.

### Statistical analysis

For the comparison of cochlear length and hair cell counts, we performed either a two-tailed unpaired Student’s t-test, or a one-way ANOVA followed by Tukey’s or Šídák’s multiple comparisons test. The Ki67- or EdU-positive ratios were evaluated using a two-tailed unpaired Student’s t-test. Student’s t-test and ANOVA were conducted using GraphPad Prism version 10 (GraphPad Software), with statistical significance set at P < 0.05. The N values indicate the number of hair cells in circular histograms and the number of mice in other experiments.

### Hair cell orientation analysis

The data comprised 11 individual experiment groups, each comparing the angle for hair bundle orientation between control and cKO mice. Measurements were taken in a patterned way for each mouse in each experiment group: from hair cells from four different rows (IHC, OHC1, OHC2, and OHC3 rows) and at two cochlear turns (the basal and middle turns), leading to 8 possible combinations.

A Bayesian mixed model analysis was run separately for each conditional KO using the R package bpnreg v2.0.3^80^ (Cremers, Jolien. n.d. Bpnreg: Bayesian Projected Normal Regression Models for Circular Data. https://github.com/joliencremers/bpnreg) with fixed effects treatment, i.e. control or conditional KO mice, and for cochlear turn (basal and middle), and hair cells from four different rows (IHC, OHC1, OHC2, and OHC3 rows) along with random effects for the individual mice. Due to the limited number of mice (n=6-8), the regression model was either a simple additive model or one containing the interaction between cochlear turn and hair cell row. The bpnreg package respects the circular nature of the angle measurements and yields samples from the posterior distributions for the regression coefficients. Convergence of the underlying Markov Chain was assessed via traceplots. The final model chosen for analysis (additive vs. interaction) was based on Watanabe-Akaike criterion. Posterior samples from the selected model were used to assess the difference in angles for each of the eight cochlear turn and hair cell row combination between conditional KO and control groups. Each difference was judged to be significant based on whether the posterior 95% credible intervals missed the origin or not (Supplementary Table 1). The circular variance ratio was defined as the circular variance of hair cell orientation in the KO group divided by the circular variance of hair cell orientation in the control group.

## Acknowledgments

We thank our lab members for insightful comments on the manuscript, W. Dong for excellent technical support, E. Fuchs (Rockefeller University) for sharing the Celsr1 antibody. The *Wnt7a^flox/flox^* mice were kindly shared by Jeremy Nathans (Johns Hopkins University), *Emx2^Cre/+^* mice by Angelika Doetzlhofer (Johns Hopkins University), and *Dermo1(Twist2)^Cre/+^* mice by Sung-Ho Huh (University of Mississippi). This work was supported by JSPS (Japan Society for the Promotion of Science) Overseas Research Fellowships (IK), NIH (National Institutes of Health) UM1TR004921 (BN), K08DC019683, R21DC022058 (TAJ), NIH/NIDCD Division of Intramural Research fund DC000094-01 (RH), R01DC021110, R01DC016919 (AGC), the Stanford Initiative to Cure Hearing Loss. This research was supported in part by the Intramural Research Program of the NIH. The contributions of the NIH authors are considered Works of the United States Government. The findings and conclusions presented in this paper are those of the authors and do not necessarily reflect the views of the NIH or the U.S. Department of Health and Human Services.

## Author contribution

I.K., E.L.S., W.D., and S.E.B. performed experiments and collected data; A.P.D., K.P.R., R.H., and T.A.J. performed computational analysis on single-cell data, B.N. and B.E., and A.P.D. performed statistical analysis on hair cell orientation, I.K., S.E.B., and A.G.C. designed experiments; I.K., A.P.D., K.P.R., B.N., and A.G.C. drafted the manuscript.

## Code Availability

The following publicly available algorithms and software were used for data analysis presented in this manuscript: Seurat (v5.3.0), CellChat (v2.1.2), and R (v4.5.2) package bpnreg (v2.0.3).

## Figure legend

**Supplementary Figure 1. In situ hybridization and hair bundle orientation in the Emx2-Wls cKO cochlea. a-b.** Negative (DapB) and positive (Polr2a) controls for *in situ* hybridization in E16.5 cochleae. **c-h.** RNA scope *in situ* hybridization of Wls mRNA in the basal, middle, and apical turn of the control (c-c’’, e-e’’, g-g’’) and Emx2-Wls cKO (d-d’’, f-f’’, h-h’’) cochleae. Magnified views (white boxes) of the cochlear epithelium and POM are shown in (c’-h’) and (c’’-h’’), respectively. In Emx2-Wls cKO cochleae, Wls mRNA is almost undetectable in the cochlear epithelium, while that in the POM remain comparable to those in control cochleae. **i-j.** Whole mount preparation of the Emx2-Wls cKO and control cochlea stained for f-actin. In the Emx2-Wls cKO cochlea, hair cells exhibit more variable orientations compared to controls. **k.** Circular histograms depicting the distribution of hair bundle orientation both control and Emx2-Wls cKO cochleae, showing significantly large variance and the average rotation for the IHC, OHC1, and OHC3 rows in the Emx2-Wls cKO cochleae. CD, cochlear duct; ST, scala tympani; SV, scala vestibule; SG, spiral ganglion. Scale bar: 20 µm in (a-h’’), 5 µm in (I, j). n, number of hair cells measured. Circular mean and variance of hair cell orientation are indicated with blue and red lines in circular histograms, respectively. A Bayesian mixed model analysis used for comparison of circular mean of hair cell orientation between control and cKO groups. *significantly different (for hair cell orientation analysis), ns = not significant.

**Supplementary Figure 2. Expression of Wnt5a, Wnt7a, and Wnt7b mRNA in the control and conditional knockout cochlea. a-d.** *Wnt5a* mRNA is robustly expressed in the epithelium and periotic mesenchyme of control cochlea (a, c) and is almost undetectable in the epithelium of Emx2-Wnt5a cKO (b, d) cochleae. **e-h.** Wnt7a mRNA is robustly expressed in the epithelium of the control cochlea (e, g) and is barely detectable in the Emx2-Wnt7a cKO (f, h) cochleae. **i-l.** Wnt7b mRNA is robustly expressed in the epithelium of the control cochlear (I, k) and is almost undetectable in the Emx2-Wnt7b cKO (j, l) cochleae. Scale bar: 20 µm.

**Supplementary Figure 3. Cochlear length and core PCP protein expression in Wnt5a, Wnt7a, and Wnt7b cKO cochlea. a-b, f-g, k-m.** Comparable lengths between Myo7a-marked controls and cochleae from Emx2-Wnt5a (a), Emx2-Wnt7a (f), and Emx2-Wnt7b cKO (k) mice at E18.5. **c, h, m** Hair cell counts show no significant differences between Emx2-Wnt5a (b) and Emx2-Wnt7a cKO (g) mice and their littermate controls, whereas those from Emx2-Wnt7b cKO (l) mice are slightly reduced compared to controls at E18.5. **d-e’’’, i-j’’’, n-o’’’** Polarized expression of Fzd6 and Dvl2 (white arrows) at the medial and lateral poles of OHCs, respectively, in control cochleae and cKO cochleae for Wnt5a (d’’’ and e’’’,), Wnt7a (i’’’ and j’’’), and Wnt7b (n’’’ and o’’’). Data are presented as mean ± S.D. and compared using an unpaired t-test. Scale bars: 200 µm in (a, f, k); 5 µm in (d-d’’, e-e’’, i-i”, j-j’’, n-n”, o-o”); 2 µm in (d’’’, e’’’, i’’’, j’’’, n’’’, o’’’). Data are presented as mean ± S.D. and compared using an unpaired t-test. *p < 0.05, ns = not significant.

**Supplementary Figure 4. Hair bundle orientation of the cochlea from Emx2-Wnt5a, Emx2-Wnt7a, and Emx2-Wnt7b cKO mice. a-d, g-j, m-p.** F-actin-stained cochleae from Emx2-Wnt5a (a-d), Emx2-Wnt7a (g-j), and Emx2-Wnt7b cKO (m-p) mice at E18.5. **e, f, k, l, q, r.** Circular histograms demonstrating no significant variance or average deviation hair bundle orientation in Emx2-Wnt5a (e, f) and Emx2-Wnt7a (k, l) cKO cochleae. and For Emx2-Wnt7b cKO (q, r) cochlea, there was no significant variance, but significantly different circular means of hair cell orientation compared to controls was detected for the IHC, OHC1, and OHC2 rows. Scale bar: 5 µm. n, number of hair cells measured. Circular mean and variance of hair cell orientation are indicated with blue and red lines in circular histograms, respectively. A Bayesian mixed model analysis used for comparison of circular mean of hair cell orientation between control and cKO groups. *significantly different (for hair cell orientation analysis), ns = not significant.

**Supplementary Figure 5. Cochlear length and core PCP protein expression after deletion of Wnts. a-b, f-g, k-l.** Whole mount preparation of cochleae from Emx2-Wnt5a; Wnt7a (a-b), Emx2-Wnt5a; Wnt7b (f-g), and Emx2-Wnt7a; Wnt7b cKO (k-l) mice and their littermate controls at E18.5. The lengths of Emx2-Wnt5a; Wnt7b and Emx2-Wnt7a; Wnt7b cKO, but not Emx2-Wnt5a; Wnt7a cKO, cochleae were significantly shorter than those of controls. **c, h, m** Hair cell counts show no significant differences between Emx2-Wnt5a; Wnt7a (c) and Emx2-Wnt7a; Wnt7b cKO (m) cochleae and their littermate controls, whereas those from Emx2-Wnt5a; Wnt7b cKO (h) cochleae are slightly reduced compared to controls at E18.5. **d-e’’’, i-j’’’, n-o’’’** Polarized expression of Fzd6 and Dvl2 is observed at the medial and lateral poles of OHCs, respectively, in control cochlea and cKO cochleae for Wnt5a and Wnt7a (d’’’ and e’’’, white arrows), Wnt5a and Wnt7b (i’’’ and j’’’, white arrows), and Wnt7a and Wnt7b (n’’’ and o’’’, white arrows). Data are presented as mean ± S.D. and compared using an unpaired t-test. Scale bars: 200 µm in (a, f, k); 5 µm in (d-d’’, e-e’’, i-i”, j-j’’, n-n”, o-o”); 2 µm in (d’’’, e’’’, i’’’, j’’’, n’’’, o’’’). **p < 0.01, ***p < 0.001, ****p < 0.0001, ns = not significant.

**Supplementary Figure 6. Hair bundle orientation after deletion of Wnt members the cochlear epithelium. a-d, g-j, m-p.** F-actin-stained cochleae from Emx2-Wnt5a; Wnt7a (a-d), Emx2-Wnt5a; Wnt7b (g-j), and Emx2-Wnt7a; Wnt7b cKO (m-p) mice at E18.5. **e, f, k, l, q, r.** Circular histograms demonstrating the distribution of hair bundle orientation in the cochlea from Emx2-Wnt5a; Wnt7a (e, f), Emx2-Wnt5a; Wnt7b (k, l), and Emx2-Wnt7a; Wnt7b cKO (q, r) mice at E18.5. Emx2-Wnt5a; Wnt7a cKO cochleae show normal hair cell orientation. Emx2-Wnt5a; Wnt7b cKO cochleae exhibit more variable orientations of hair cells compared to controls (g-j), confirmed by significantly larger circular variance in hair cell orientation, but the average orientation was not significantly different from controls, with the exception of the IHC row of the basal turn. Similarly, hair cell variance in Emx2-Wnt7a; Wnt7b cKO cochleae was not significantly different from controls, with no significantly different circular mean except for the OHC3 row. Scale bar: 5 µm. n, number of hair cells measured. Circular mean and variance of hair cell orientation are indicated with blue and red lines in circular histograms, respectively. A Bayesian mixed model analysis used for comparison of circular mean of hair cell orientation between control and cKO groups. *significantly different, ns = not significant.

**Supplementary Figure 7. Hair bundle orientation after deletion of three Wnt members in the cochlear epithelium.** Whole mount preparation of f-actin-stained cochlea from control (a, a’) and Emx2-Wn5a; Wnt7a; Wnt7b cKO (b, b’) mice. **i** Circular histograms illustrating the distribution of hair bundle orientations in the middle turn of both control and Emx2-Wn5a; Wnt7a; Wnt7b cKO cochleae, showing no significant difference in circular means and variance from controls. n, number of hair cells measured. Circular mean and variance of hair cell orientation are indicated with blue and red lines in circular histograms, respectively. A Bayesian mixed model analysis used for comparison of circular mean of hair cell orientation between control and cKO groups. ns = not significant.

**Supplementary Figure 8. In situ hybridization of Wnt5a mRNA in the Dermo1-Wnt5a cKO cochlea.** Wnt5a mRNA is robustly expressed in the cochlear duct and periotic mesenchyme of the control cochleae (*Dermo1^Cre/+^; Wnt5a^flox/+^*) (a-a’, c-c’), and notably absent in the periotic mesenchyme of *Dermo1^Cre/+^; Wnt5a^flox/flox^* cochleae (Dermo1-Wnt5a) (b-b’, d-d’) at E16.5. Wnt5a mRNA remain present in the cochlear epithelium and spiral ganglion (b-b’, d-d’), indicating effective deletion of Wnt5a mRNA specifically from the POM. **e-g.** F-actin-stained middle turn of cochleae from control (e) and Dermo1-Wnt5a cKO (f) embryos at E18.5. **i** Circular histograms demonstrating comparable hair bundle orientation in the middle turn of control and Dermo1-Wnt5a cKO cochleae. CD cochlear duct; ST scala tympani; SV scala vestibule; SG spiral ganglion. Scale bar: 20 µm in (a-h’’), 5 µm in (I, j). A Bayesian mixed model analysis used for comparison of circular mean of hair cell orientation between control and cKO groups. ns = not significant.

**Supplementary Figure 9. Deleting Wntless in the cochlear epithelium and periotic mesenchyme at E11.5/12.5 causes mild PCP defects. a.** Schematic of broadly deleting Wntless using the Sox9-CreERT2; Wntless-flox mice. Tamoxifen (0.2 mg/g) was administered at E11.5 and E12.5 and cochleae were analyzed at E18.5. **b.** Sox9-Wls cKO cochlea is significantly shorter than controls. **c-d.** Length and hair cell counts of Sox9-Wls cKO cochleae with tamoxifen administered at E11.5-12.5 were significantly lower than controls, but higher than Sox9-Wls cKO cochleae with tamoxifen administered at E10.5-11.5. **e-h.** Whole mount preparation of f-actin-stained cochlea showing hair cells malrotated towards the apex in the Sox9-Wls cKO cochlea (arrows). **i-j.** Circular histograms demonstrating the distribution of hair bundle orientation in control and Sox9-Wls cKO cochleae. Significantly larger circular variance and different circular means of hair cell orientation are detected in the basal and middle turns of Sox9-Wls cKO cochleae. **k-s.** Control and Sox9-Wls cKO cochleae showed polarized expression of core PCP proteins in (Fzd6 (k-l), Celsr1 (m-n), Vangl1 (p-q), and Vangl2 (r-s)). Scale bar: 200 µm in (b), 5 µm in (e-h, k-s). Data are presented as mean ± S.D. and analyzed using one-way ANOVA followed by Tukey’s multiple comparisons test. n, number of hair cells measured. Circular mean and variance of hair cell orientation are indicated with blue and red lines in circular histograms, respectively. A Bayesian mixed model analysis used for comparison of circular mean of hair cell orientation between control and cKO groups. *significantly different (for hair cell orientation analysis), **p < 0.01, ***p < 0.001, ****p < 0.0001, ns = not significant.

**Supplementary Figure 10. Loss of Wls mRNA expression in Sox9-Wls cKO cochleae. a-f’** In situ hybridization showing robust Wls mRNA expression in all turns of control (a-a’, c-c’, e-e’) cochleae, and loss of expression in Sox9-Wls cKO (b-b’, d-d’, f-f’) cochleae at E18.5. Scale bar: 50 µm. CD cochlear duct; ST scala tympani; SV scala vestibule; SG spiral ganglion.

**Supplementary Figure 11. Hair bundle orientation and proliferation in Sox9-Wls cochlea after Wntless at E10.5 and E11.5. a-b’.** F-actin-stained cochlea from Sox9-Wls cKO (b, b’) embryos showing five to six rows of hair cells, disorganized hair cells lacking a clear separation between IHCs and OHCs, and malrotation of hair cells toward the apex. **c.** Circular histograms illustrating the distribution of hair bundle orientation in IHCs and OHCs in the middle turn of control and Sox9-Wls cKO cochleae. Sox9-Wls cKO cochleae have significantly larger circular variance for combined IHC and OHCs and significantly different circular means of hair cell orientation for each of IHC and OHC rows compared to controls. **d-e.** Whole-mount preparation of E15.5 cochlea labeled with EdU or immunostained for Ki67, showing comparable degree of proliferation of Sox2-positive cells in control and Sox9-Wls cKO organs. **f-g.** Proliferation of POM cells in the POM was not significantly different between control and Sox9-Wls cKO cochleae at E15.5. Data are presented as mean ± S.D. and analyzed using an unpaired t-test (d-e). n, number of hair cells measured. Circular mean and variance of hair cell orientation are indicated with blue and red lines in circular histograms, respectively. A Bayesian mixed model analysis used for comparison of circular mean of hair cell orientation between control and cKO groups. *significantly different (for hair cell orientation analysis), ns = not significant.

