## Supplementary Figures for "Epithelial-Mesenchymal Wnt Crosstalk Directs Planar Cell Polarity in the Developing Cochlea"

Supplementary Figure 1

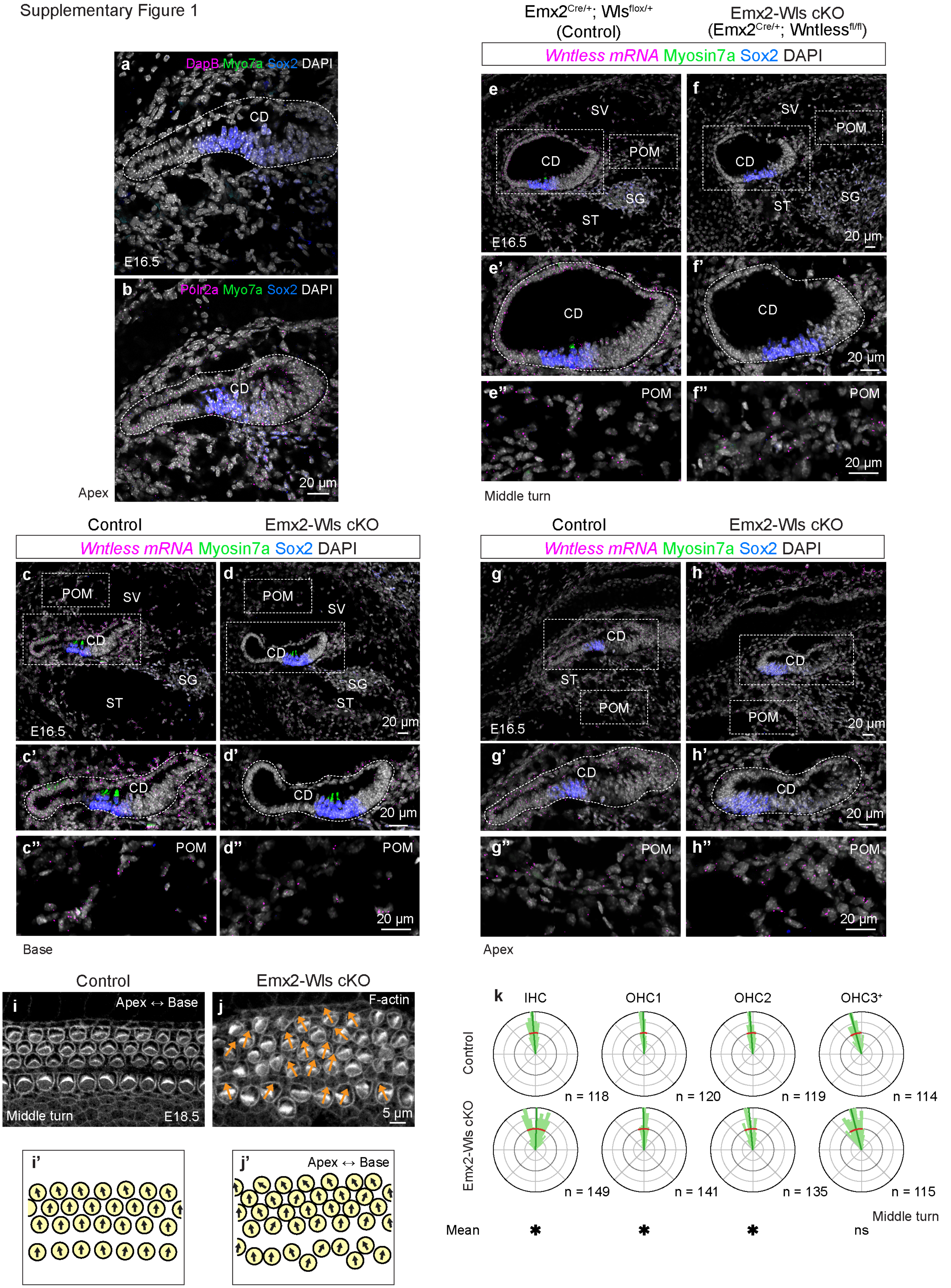

Supplementary Figure 2

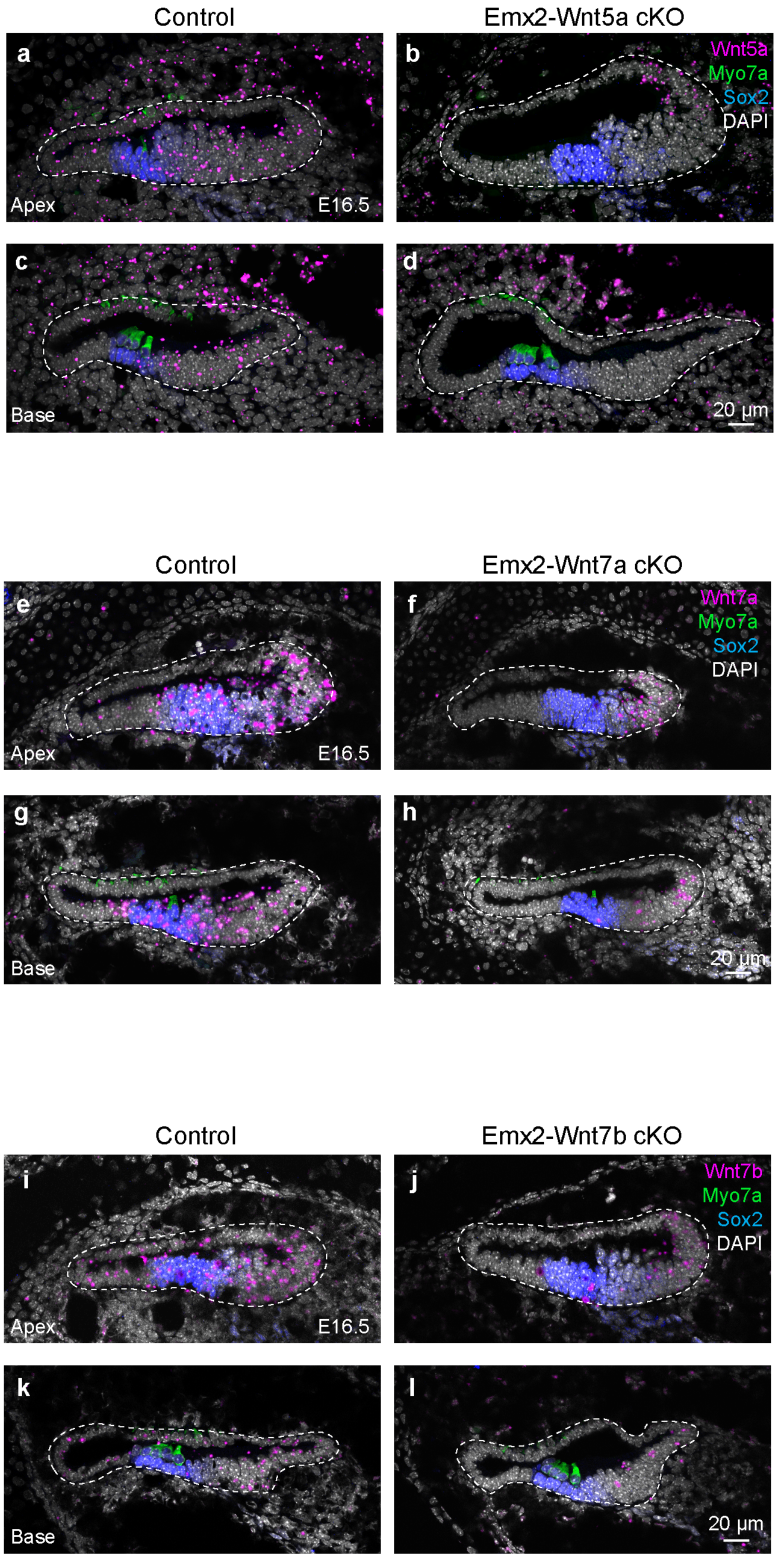

Supplementary Figure 3

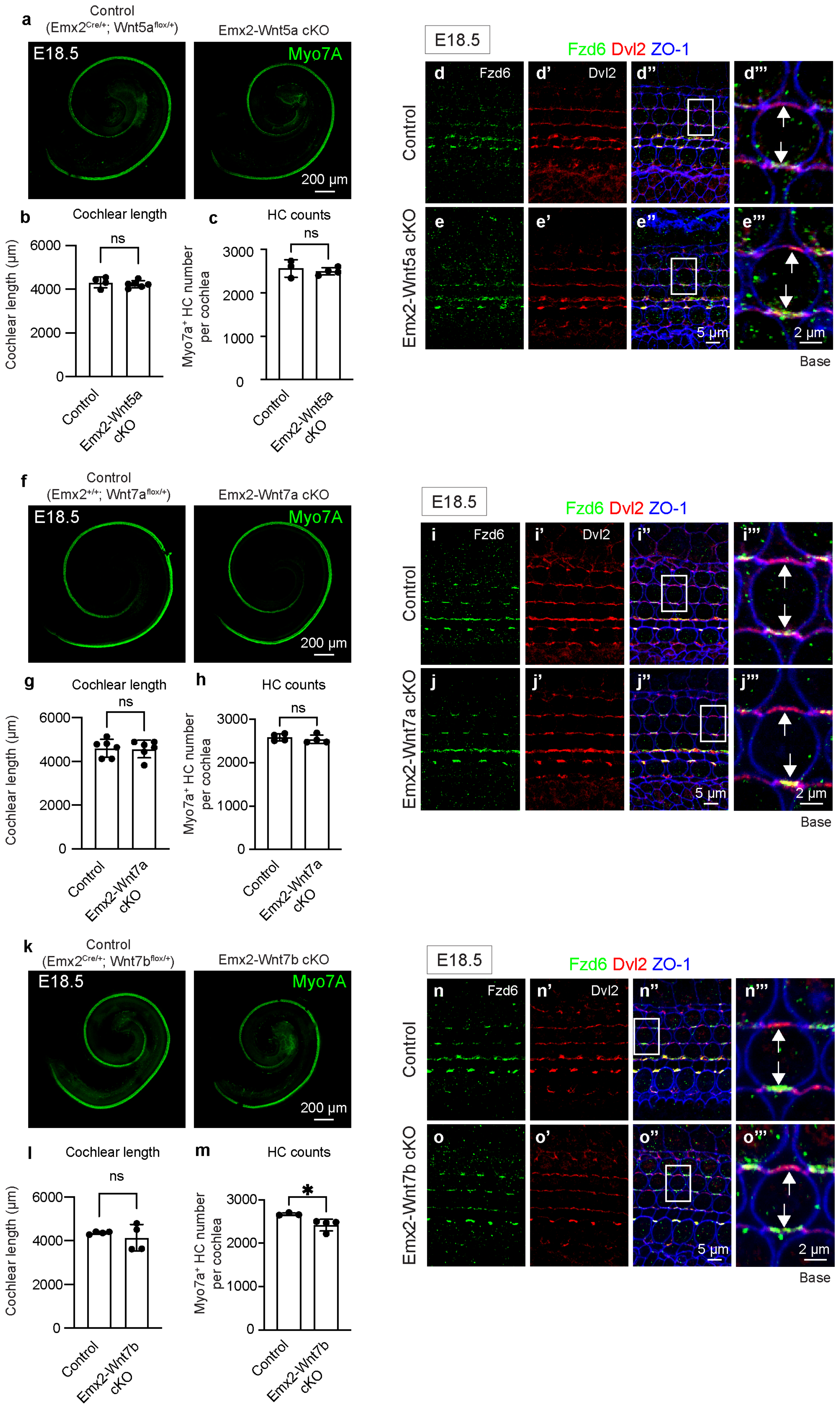

Supplementary Figure 4

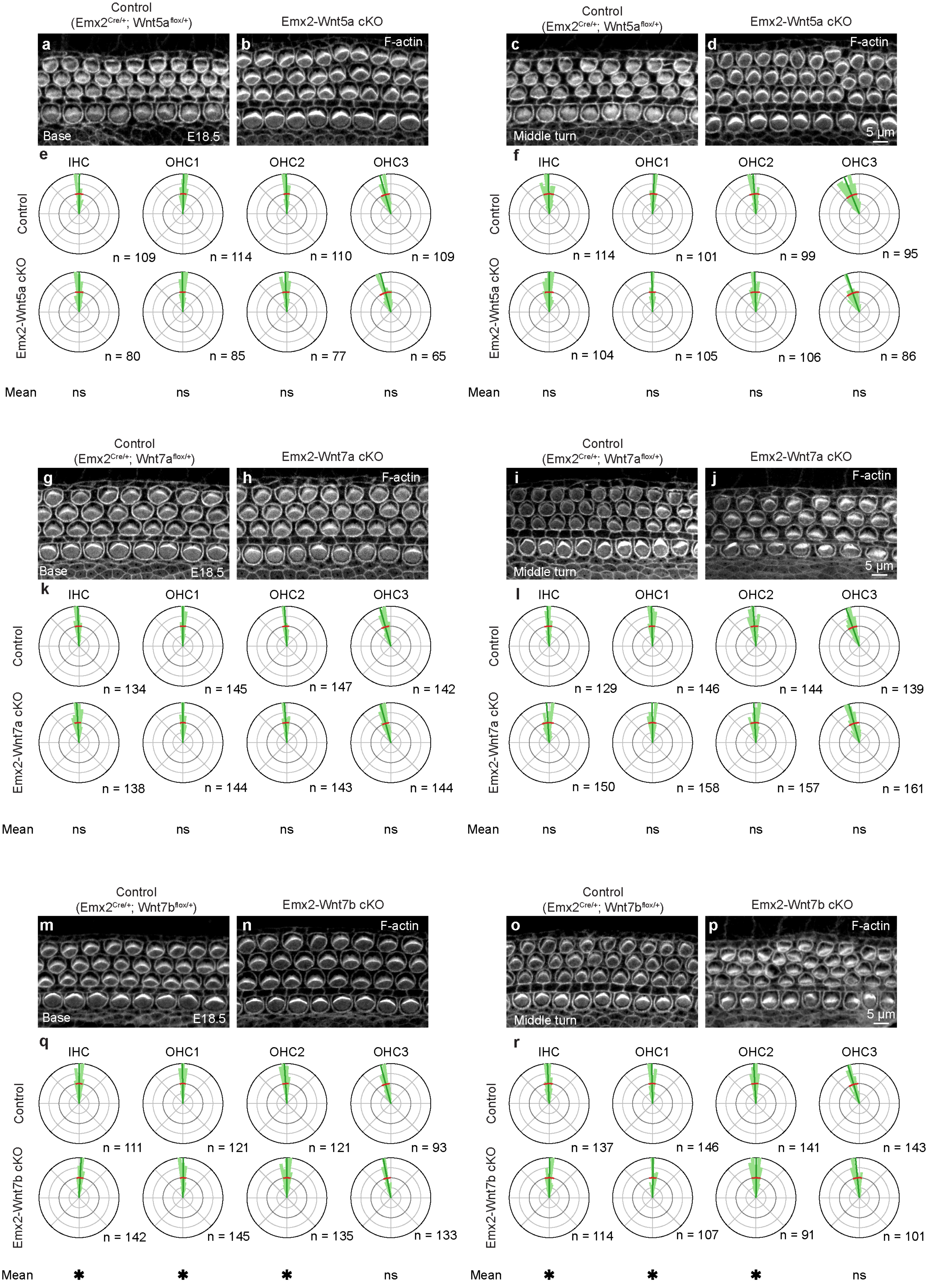

Supplementary Figure 5

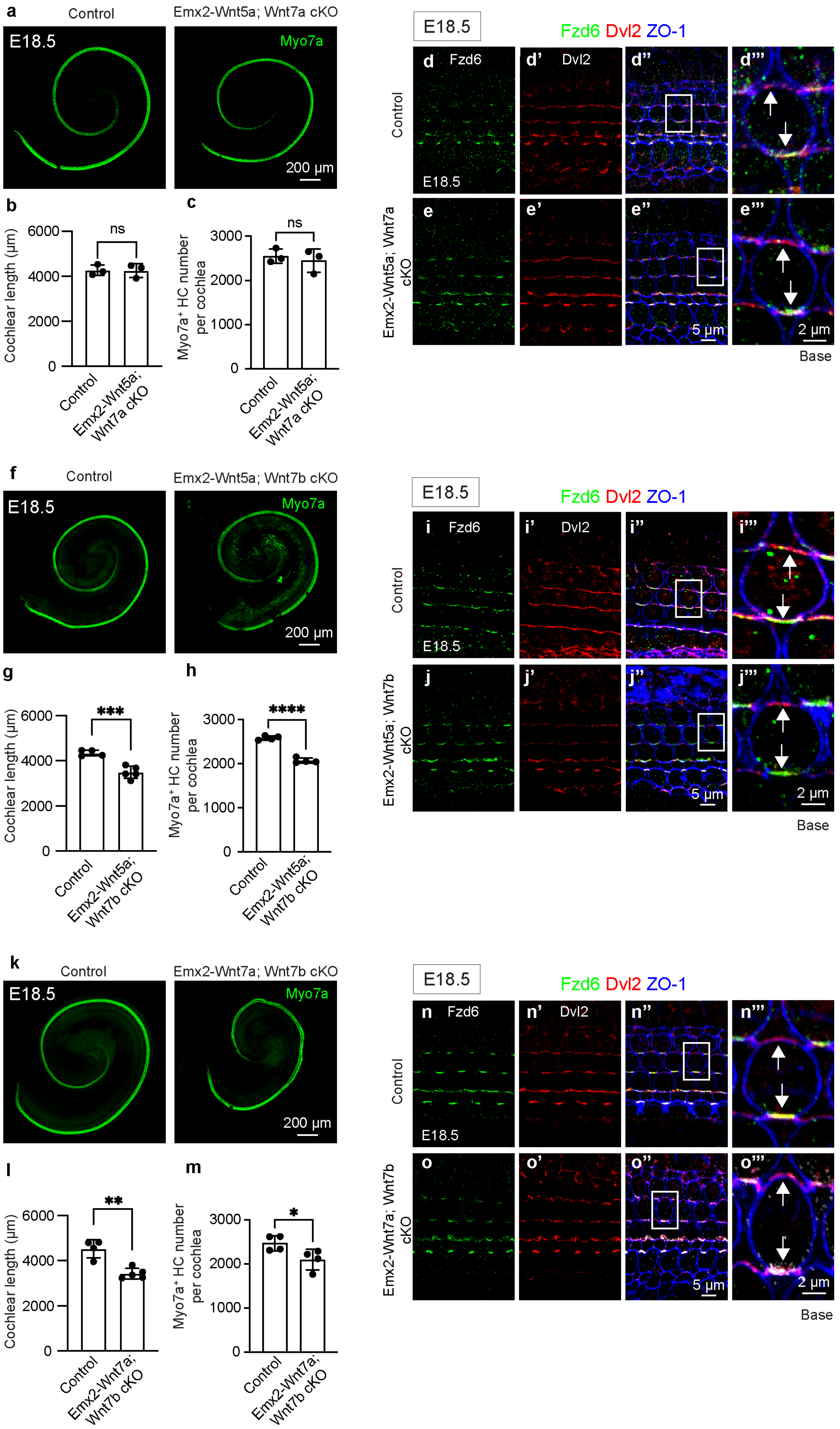

Supplementary Figure 6

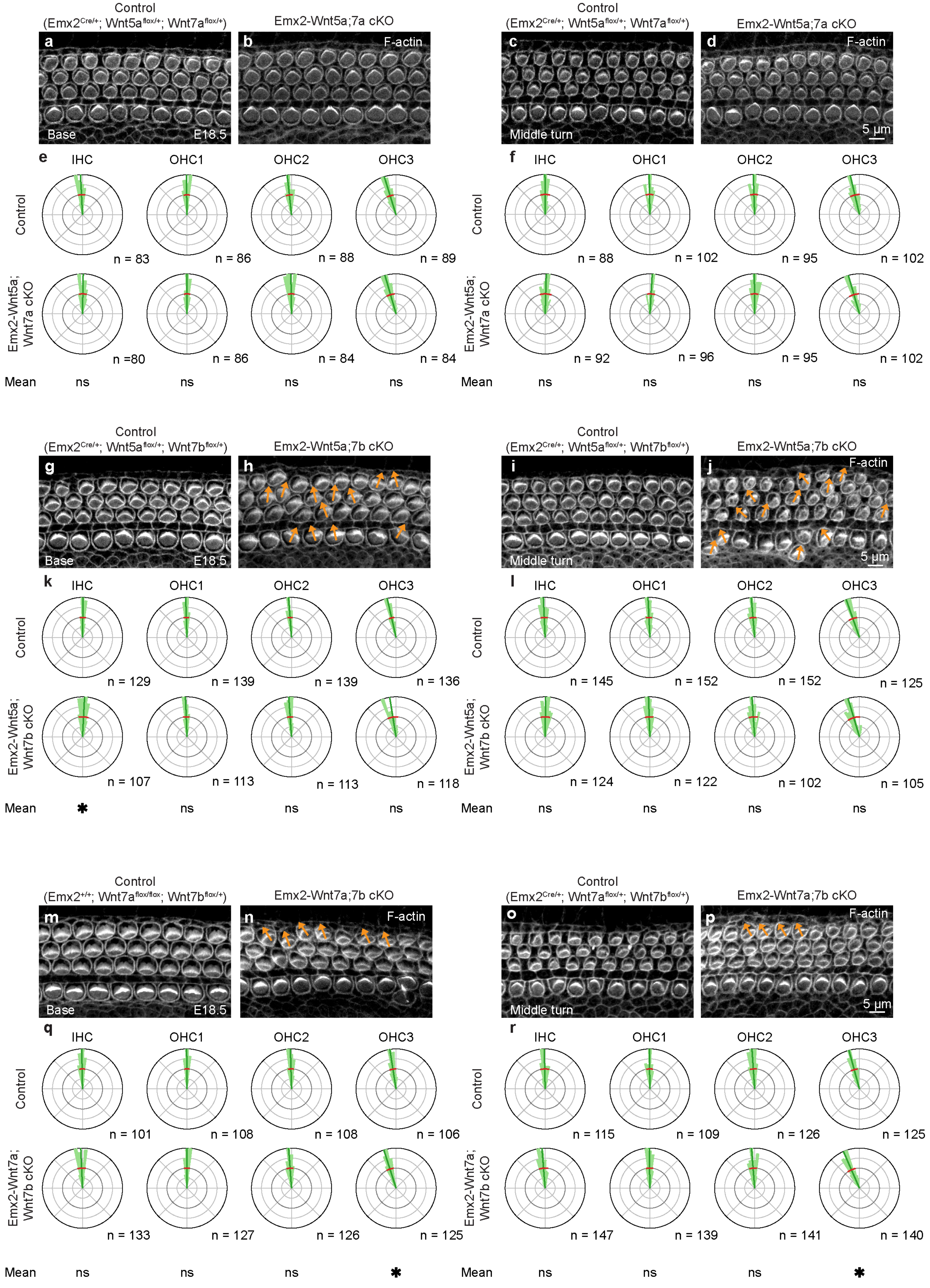

Supplementary Figure 7

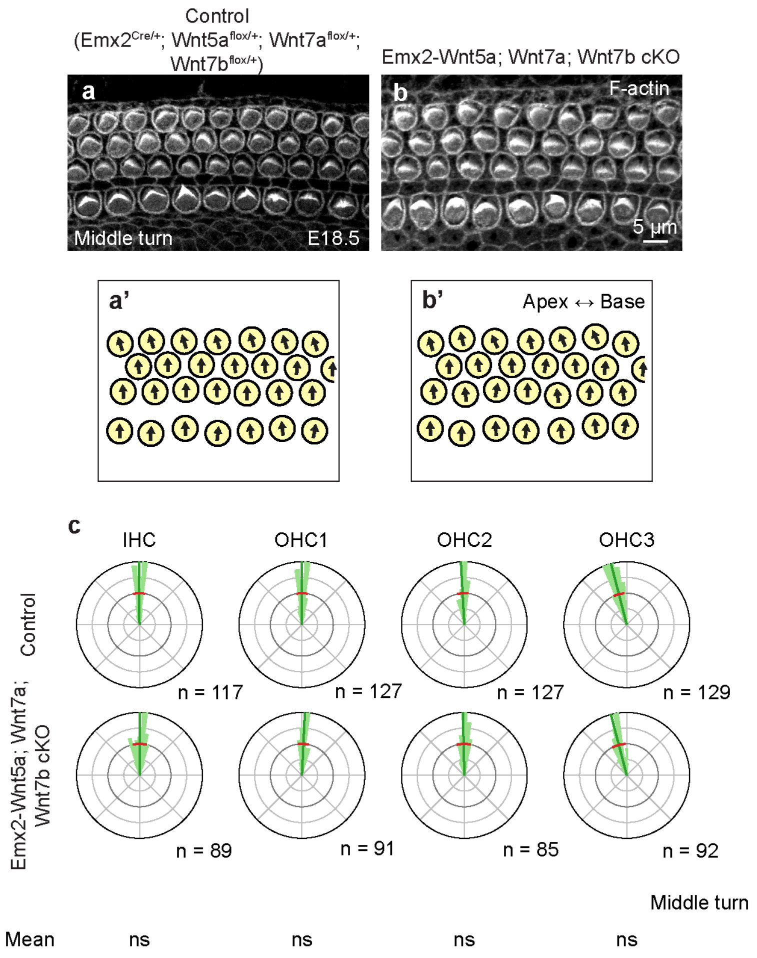

Supplementary Figure 8

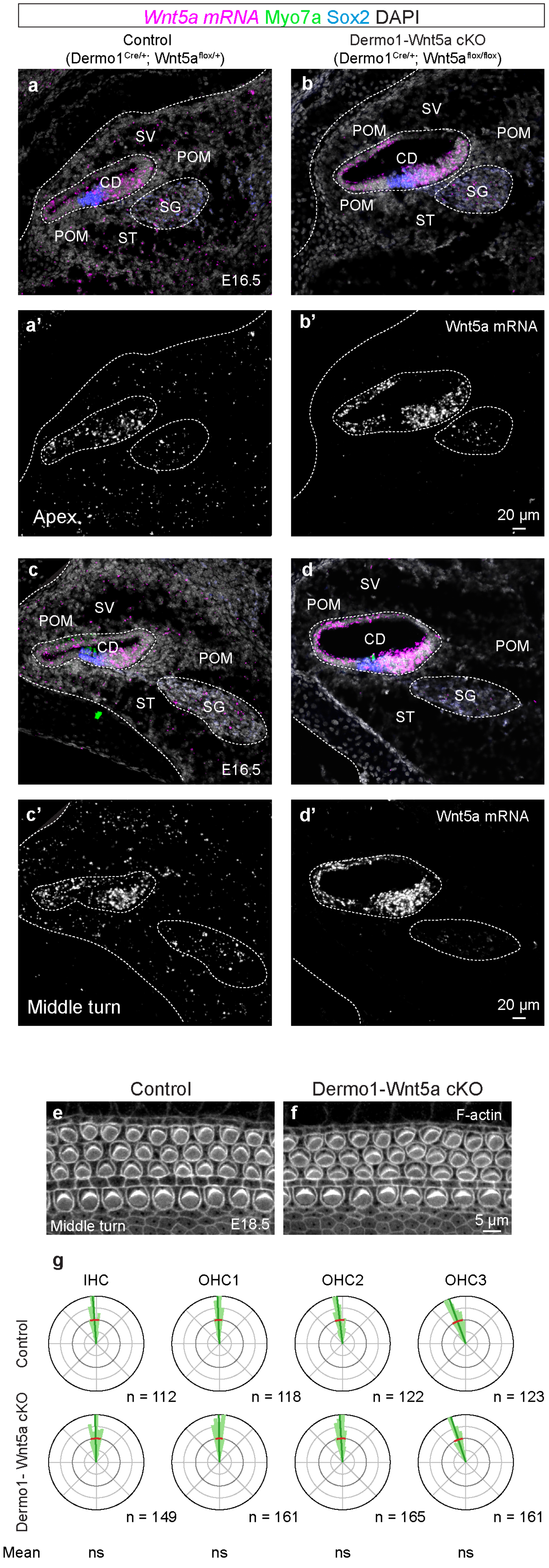

Supplementary Figure 9

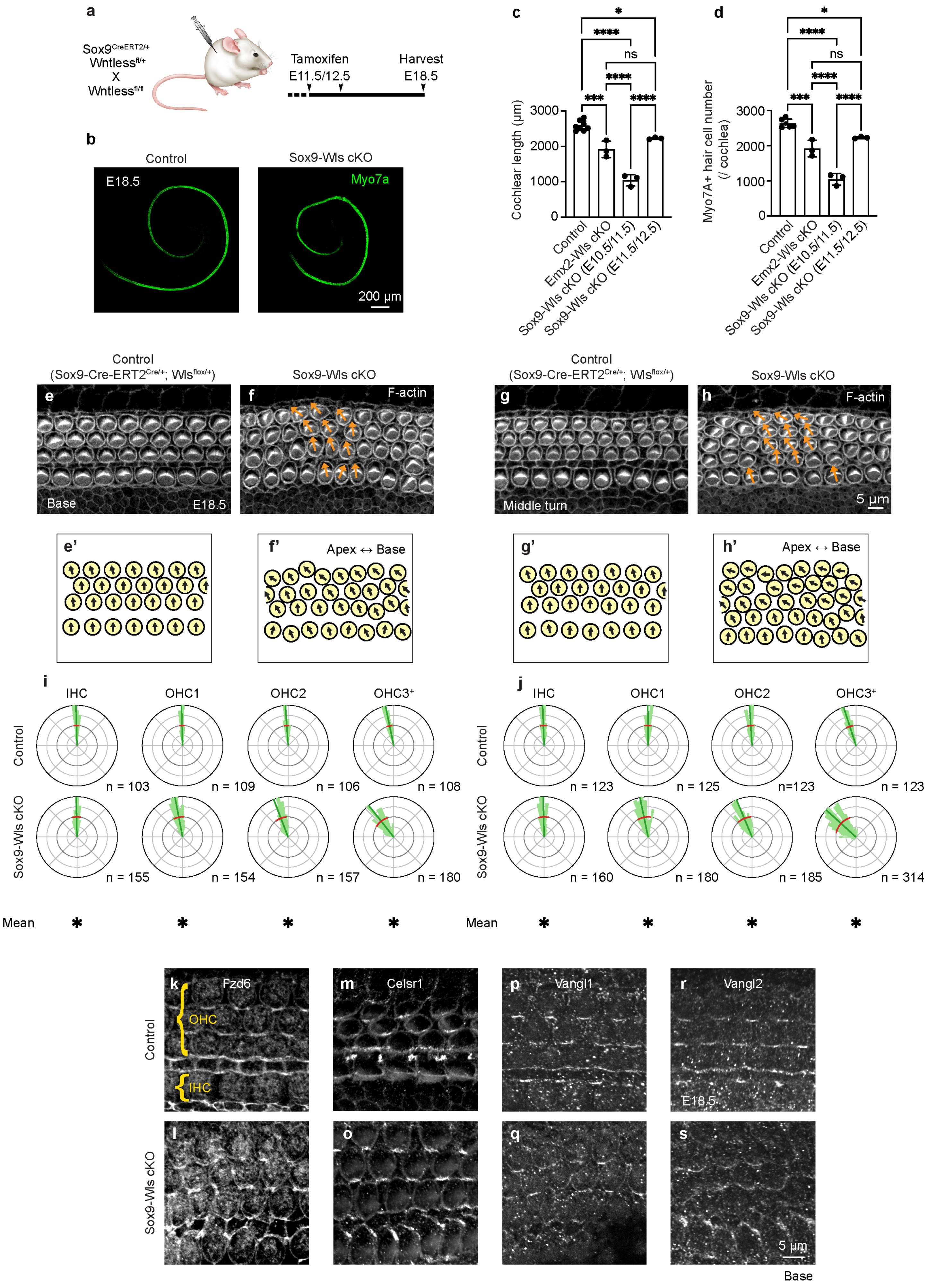

Supplementary Figure 10

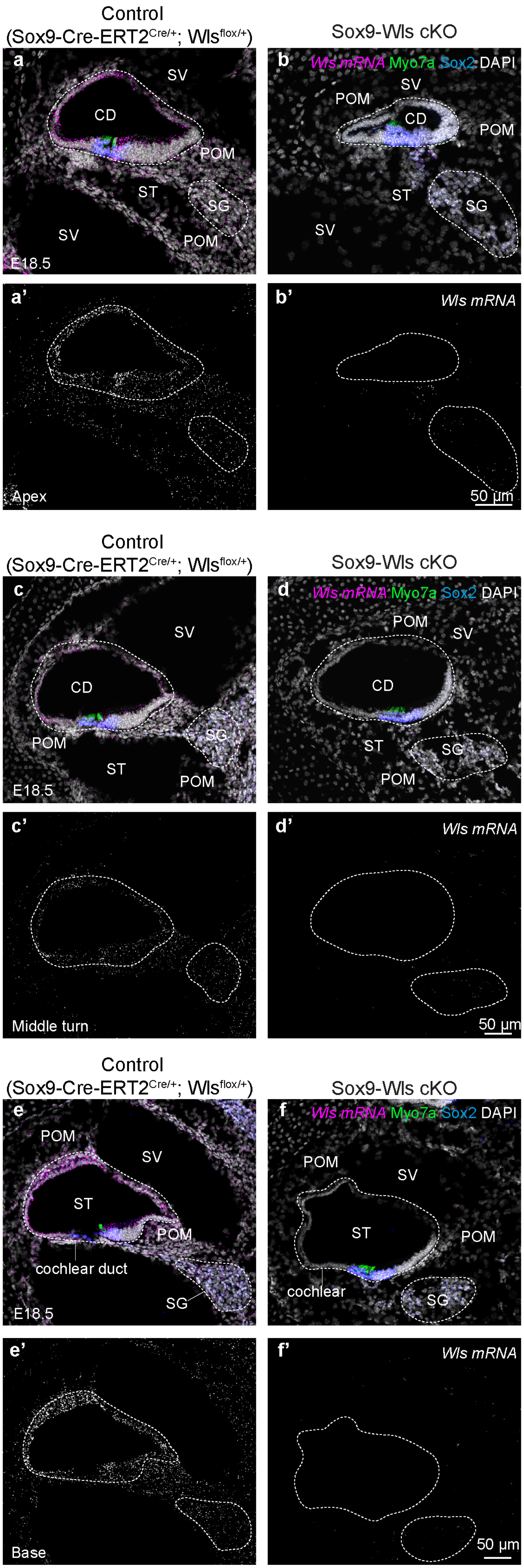

Supplementary Figure 11

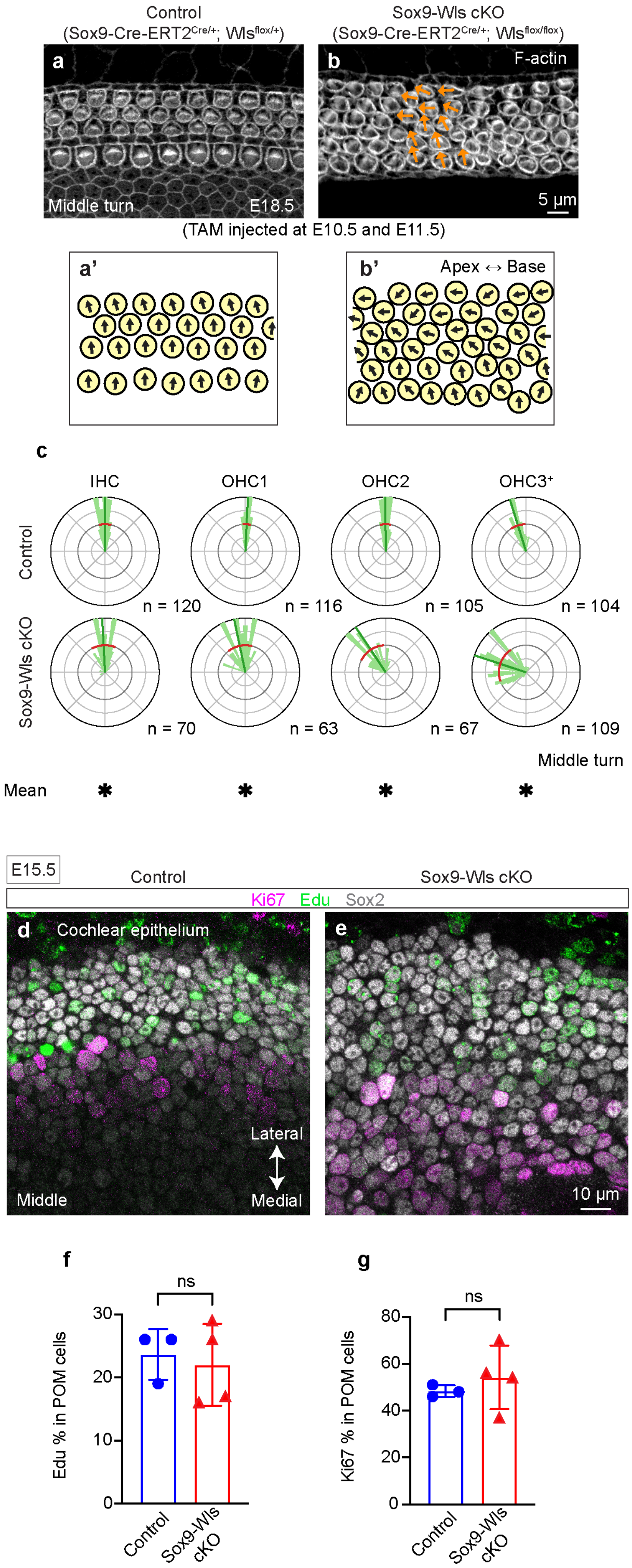
